# Cellular and genetic drivers of RNA editing variation in the human brain

**DOI:** 10.1101/2021.07.16.452690

**Authors:** Ryn Cuddleston, Junhao Li, Xuanjia Fan, Alexey Kozenkov, Matthew Lalli, Shahrukh Khalique, Stella Dracheva, Eran A. Mukamel, Michael S. Breen

**Affiliations:** Department of Psychiatry at Mount Sinai, New York, New York, 10029, USA; Department of Genetics and Genomic Sciences at Mount Sinai, New York, New York, 10029, USA; Seaver Autism Center for Research and Treatment at Mount Sinai, New York, New York, 10029, USA; Pamela Sklar Division of Psychiatric Genomics at Mount Sinai, New York, New York, 10029, USA; James J Peters VA Medical Center, Bronx, New York, 10468, USA; Department of Cognitive Science, University of California, San Diego, La Jolla, CA 92037, USA

## Abstract

Posttranscriptional adenosine-to-inosine modifications amplify the functionality of RNA molecules in the brain, yet the cellular and genetic regulation of RNA editing is poorly described. We quantified base-specific RNA editing across three major cell populations from the human prefrontal cortex: glutamatergic neurons, medial ganglionic eminence GABAergic neurons, and oligodendrocytes. We found more selective editing and RNA hyper-editing in neurons relative to oligodendrocytes. The pattern of RNA editing was highly cell type-specific, with 189,229 cell type-associated sites. The cellular specificity for thousands of sites was confirmed by single nucleus RNA-sequencing. Importantly, cell type-associated sites were enriched in GTEx RNA-sequencing data, edited ∼twentyfold higher than all other sites, and variation in RNA editing was predominantly explained by neuronal proportions in bulk brain tissue. Finally, we discovered 661,791 cis-editing quantitative trait loci across thirteen brain regions, including hundreds with cell type-associated features. These data reveal an expansive repertoire of highly regulated RNA editing sites across human brain cell types and provide a resolved atlas linking cell types to editing variation and genetic regulatory effects.

## INTRODUCTION

The complexity of the central nervous system (CNS) is largely coordinated through multiple layers of transcriptional regulation, generating functionally distinct RNA molecules with specialized post-transcriptional modifications^1, 2^. Adenosine to inosine (A-to-I) RNA editing is abundant in the human brain and predicted to occur at millions of locations across the genome^3, 4^. A-to-I editing occurs at single isolated adenosines (selective editing) as well in extended regions with multiple neighboring adenosines (RNA hyper-editing)^5–7^, and is catalyzed by adenosine acting on RNA (ADAR) enzymes. These base-specific changes exponentially amplify RNA sequence diversity and expand the functionality of many brain expressed genes, by allowing the same coding sequence to produce different mRNA and products. Although the growth of RNA-sequencing data sets has increased the catalogue of known editing sites, the functional relevance of most sites remains unknown. We reasoned that sites with precisely regulated differences in RNA editing across brain cell types or brain regions signal a potentially critical role in supporting the functional diversity of brain circuits. Moreover, dissecting the genetic regulation of RNA editing at these sites gives insight into their potential role in healthy and diseased brains.

In the CNS, RNA editing regulates neuronal transcription, splicing and subcellular localization of mRNA transcripts^1, 8–1^^1^. RNA editing in protein-coding regions can result in recoding specific amino acids, which influences essential neurodevelopmental processes, including actin cytoskeletal remodeling at excitatory synapses^1,1^^2^, regulation of gating kinetics of inhibitory receptors^1, 13^, and modulation of neurotransmission at inhibitory synapses^1, 14^. RNA editing sites are dynamically regulated throughout human cortical development^15, 16^, with marked increases in editing levels occurring between mid-fetal development and infancy. These profiles are conserved in non-human primates and murine models of cortical development, indicating an evolutionarily selected function^12, 15^. Moreover, widespread changes in RNA editing are linked to several neuropsychiatric and neurodevelopmental disorders^17–22^. Yet, the cellular specificity of RNA editing sites in the human brain remains largely unknown due to the lack of cell type-specific studies.

The vast majority of RNA editing sites have been detected in bulk brain tissue, a mixture of dozens of neuronal and glial cell types with distinct transcriptional, and potentially epitranscriptional, programs. In mouse brain, editing levels are higher in neurons relative to glial cells, and these trends are consistent across brain development^23, 24^. In *Drosophila*, RNA editing has been explored across several neuronal populations, each containing a unique editing signature composed of distinct site-specific editing levels in neuronal transcripts^25^. In the human brain, the challenge of purifying specific cell populations has so far prevented broad-based analysis of cell type-specific RNA editing. Single-cell RNA-sequencing has been applied in a modest number of cells^26^ (*n^cells^*=268), indicating that highly edited sites in individual cells often go undetected by bulk brain RNA-seq. The systematic identification of bona fide, high-confidence cell type-associated RNA editing sites in the human brain is critical for understanding the scope and specificity of this layer of cell regulation.

In addition to cell-specific factors, common genetic variation has also recently emerged as an important regulator of RNA editing levels in the brain^17, 27^. Integrating paired genomic and transcriptomic data can identify RNA editing quantitative trait loci (edQTLs). It is estimated that anywhere between ∼10-30% of RNA editing sites identified in bulk tissue samples of the human brain are regulated by common genetic variants^17, 27, 28^, and this number will continue to grow with increasing sample sizes. However, there has been no investigation of cell type-associated edQTLs in the human brain.

Here we sought to expose the main cellular and genetic drivers of RNA editing variation in the human brain. We first quantified RNA editing among three major cell populations purified from the adult prefrontal cortex (PFC): medial ganglionic eminence (MGE)–derived inhibitory GABAergic interneurons (MGE-GABA), excitatory glutamatergic neurons (GLU), and oligodendrocytes (OLIG). These cell populations were isolated from nine donors using fluorescence-activated nuclei sorting (FANS), followed by transcriptomic analysis by RNA-seq^29^. By combining our high-resolution cell-type-specific data with complementary single nucleus RNA-seq, as well as bulk RNA-seq from multiple brain regions from the GTEx project, we provide comprehensive analysis and validation of cell type-specific RNA editing across the human brain, including key genetic regulators of editing quantitative trait loci (edQTLs). Overall, our study suggests that RNA editing plays a critical role in regulating the diverse molecular identities of brain cell types.

## RESULTS

### Global editing rates in MGE-GABAergic interneurons, glutamatergic neurons and oligodendrocytes

To better understand differences of RNA editing between MGE-GABA, GLU and OLIG cells, we computed an Alu Editing Index (AEI) as a global measure of site selective RNA editing activity for each cell type (*see Methods*). The AEI is defined as the ratio of the total number of A-to-G edited reads over the total coverage of all adenosines in Alu elements across the transcriptome. The AEI was higher in neurons compared to OLIG (Cohen’s d = 2.58, *p*=1.2×10^−6^, linear regression) and elevated in MGE-GABA relative to GLU (Cohen’s d = 1.46, *p*=0.009, linear regression) (**Figure 1A, Supplemental Table 1**). Given that the vast majority of RNA editing occurs in Alu elements and nearly all adenosines in Alu repeats are targeted by ADARs, we queried ADAR expression levels and identified higher expression of *ADAR1* in MGE-GABA and GLU cells relative to OLIG (Cohen’s d = 4.34, *p*=8.2×10^−11^, linear regression), higher expression of *ADAR2* in MGE-GABA neurons relative to GLU and OLIG (Cohen’s d = 1.96, *p*=0.0002, linear regression), and higher *ADAR3* expression in OLIG relative to neurons (Cohen’s d = 1.78, *p*=0.001, linear regression) (**Figure 1B**). Notably, variation in the AEI was positively associated with *ADAR2* (*R*^2^=0.68) and *ADAR1* (*R*^2^=0.64) expression and negatively associated with *ADAR3* (*R*^2^=-0.33) (**Figure 1C**).

**Figure 1.**
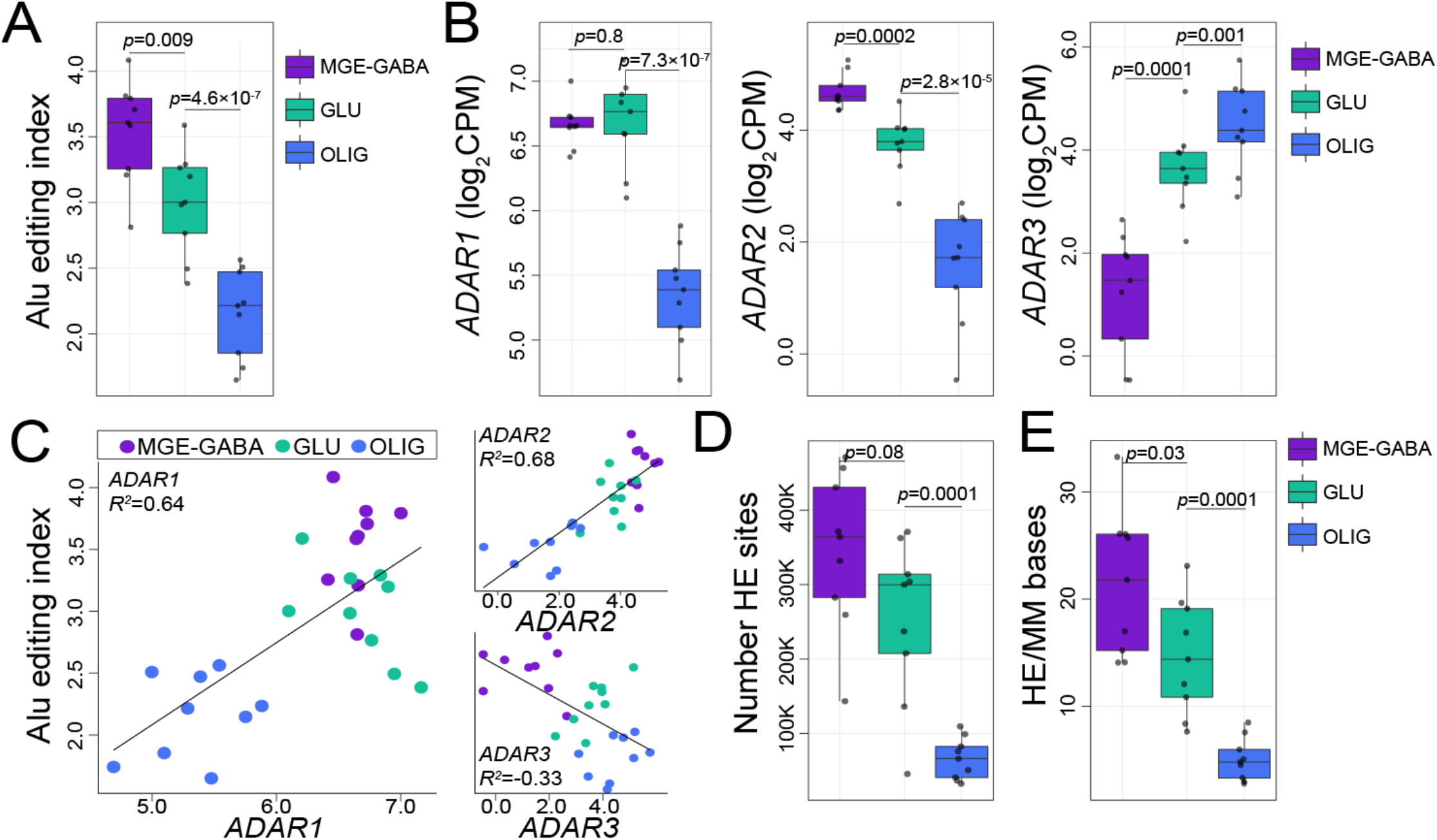
Global selective editing and hyper-editing in purified cortical cell types. **(A)** The Alu editing index (AEI) (y-axis) and **(B)** expression profiles for ADAR1, ADAR2, and ADAR3 (y-axis) were computed for each FANS-derived cell population from autopsied human adult prefrontal cortex (PFC, n=9): MGE-GABA, GLU, and OLIG populations. **(C)** The amount of AEI variance explained (R^2^, y-axis) by ADAR expression was quantified. **(D)** The total number of hyper-editing sites (y-axis) and **(E)** normalized hyper-editing rates (hyper-editing sites per million mapped bases, y-axis) were calculated across all cell types. For panels A, B, D, and E, regression analysis computed significance in mean differences between MGE-GABA vs GLU and GLU vs. OLIG. In all panels, MGE-GABA vs. OLIG is statistically significant (*p* < 0.05).

We also computed a global metric of A-to-G hyper-editing, defined as consecutive editing across many neighboring adenosines within an extended region in the same transcript, which leverages unmapped RNA reads (*see Methods*) (**Supplemental Table 1**). There were ∼4-5 times more RNA hyper-editing sites in MGE-GABA (µ=345,736 sites) and GLU (µ=253,135 sites) neurons than in OLIG (µ=66,077 sites) (Cohen’s d = 2.64, *p*=2.6×10^−6^, linear regression) (**Figure 1D**). To minimize technical variability and facilitate a direct comparison across all cell types, we normalized the hyper-editing signal to the number of mapped bases per sample and again observed a preponderance of hyper-editing in neurons relative to OLIG (Cohen’s d = 2.24, *p*=1.0×10^−5^, linear regression) and an elevated rate in MGE-GABA relative to GLU (Cohen’s d = 1.41, *p*=0.03, linear regression) (**Figure 1E**). Normalized hyper-editing rates were also positively associated with *ADAR1* (*R^2^*=0.38) and *ADAR2* (*R^2^*=0.61), but negatively associated with *ADAR3* (*R^2^*=-0.27) (**Figure S1**). Hyper-editing sites commonly occurred in introns and 3’UTRs (**Supplemental Table 1**) and enriched for a local RNA editing sequence motif whereby guanosine is depleted −1bp upstream and enriched +1bp downstream the target adenosine (**Figure S1D-E**); validating the accuracy of our hyper-editing approach. Taken together, we show that global selective editing and RNA hyper-editing are more prevalent in MGE-GABAergic than glutamatergic neurons, followed by OLIG cells. This variation in cell-specific editing activity may be due to variation in ADAR expression.

### Identification and annotation of bona fide site selective RNA editing sites

To catalogue high-confidence bona fide selective RNA editing sites, we combined *de novo* calling with a supervised approach (*see Methods*). All sites were subjected to rigorous down-stream filtering and quality control. In brief, thresholds were set to control total read coverage (>10 supporting reads), minimum edited read coverage (>3 supporting edited reads), minimum editing ratio (at least 5%). We further required that sites meet these criteria in at least 8 out of 9 donors. Sites in homopolymeric and blacklisted regions of the genome were discarded, along with sites masked as common genomic variants (**Figure 2A**). Overall, we identified a total of 107,999 sites on 20,288 genes in MGE-GABA, 109,734 sites on 19,264 genes in GLU and 64,374 sites on 11,564 genes in OLIG populations (**Supplemental Table 2**), discovery rates consistent global editing activity (**Figure 1**). We defined ∼60% of these sites as “cell type-specific” as they were uniquely detected in one cell type, and their detection rates were largely explained by cell type-specific gene expression and read coverage differences (**Figure S2**).

**Figure 2.**
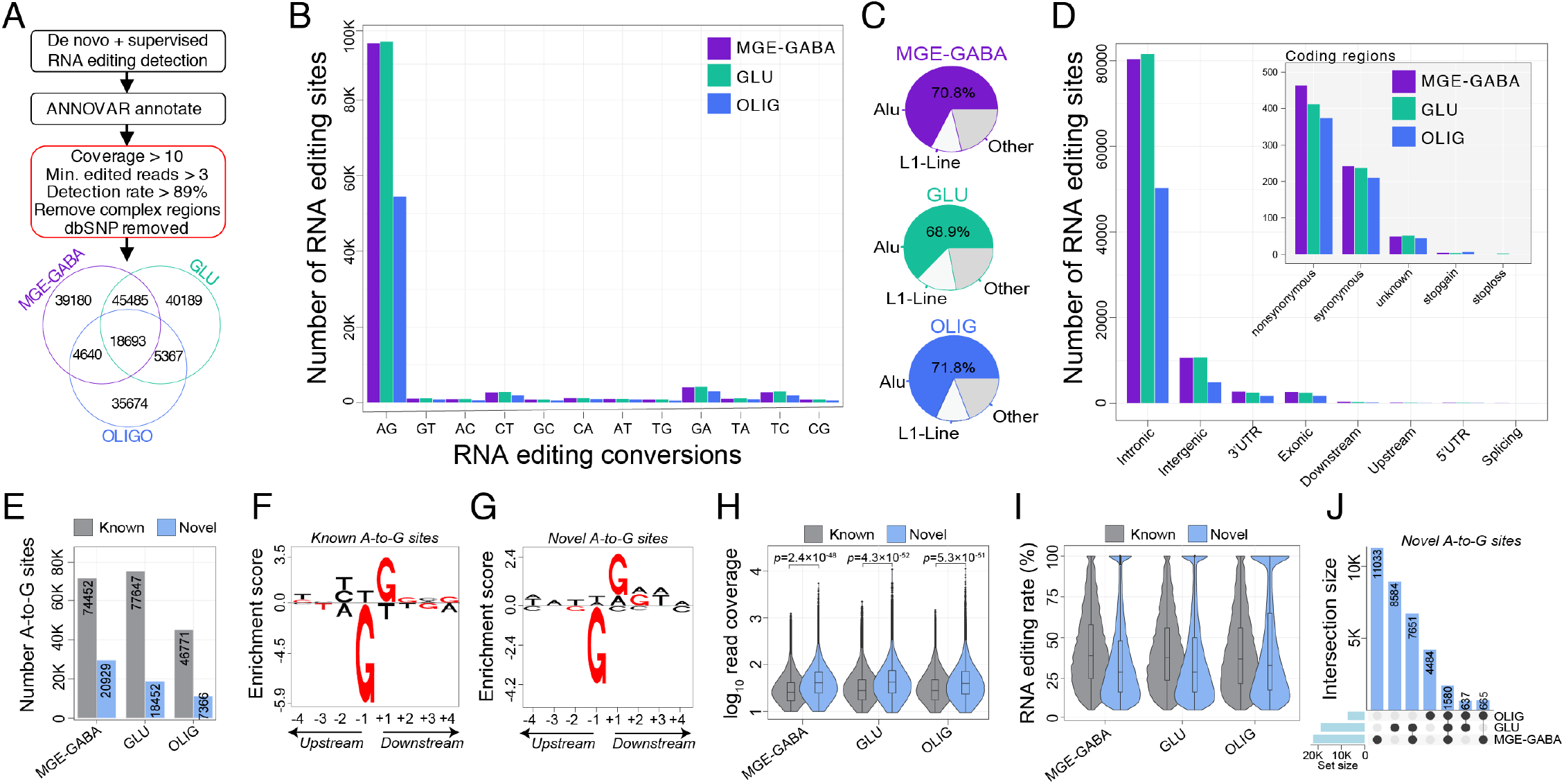
Identification and annotation of selective RNA editing sites. **(A)** Workflow for detecting high-confidence cell type-associated RNA editing sites using conjunctional *de novo* and supervised RNA editing detection approaches. The total number of sites detected per cell type are displayed in Venn diagram. **(B)** The total number of RNA editing sites by substitution class. **(C)** The fraction of RNA editing sites that map to Alu and L1-line elements relative to all other elements. **(D)** The total number of RNA editing sites by genic region. Inset plot shows the breakdown in coding regions. **(E)** The number of A-to-G editing sites (y-axis) were grouped based on novelty (known or novel) and independently numerated per cell type. Local sequence motif enrichment analysis for both **(F)** known and **(G)** novel RNA editing depicting a depletion of guanosine +1 bp upstream and enrichment of guanosine −1 bp downstream of the target adenosine. Known and novel sites for each cell population were further scrutinized by comparison of **(H)** read coverage (log_10_, y-axis) and **(I)** editing level (%, y-axis). **(J)** An upset plot highlights the cell-type convergence/divergence specifically of novel A-to-G sites.

To validate the accuracy of our approach, we annotated all events and observed consistent hallmarks of RNA editing. First, the vast majority of sites were A-to-G edits (∼86%) (**Figure 2B**) and resided within Alu elements (∼68%) (**Figure 2C**). Second, ∼73% of editing events were detected in introns, while only a small fraction impacted protein coding regions (∼0.67%) (**Figure 2D**). Third, while most RNA editing sites were known events documented in the REDIportal database (**Figure 2E**), we also identified thousands of novel A-to-G events, including 20,929 sites in MGE-GABA, 18,452 sites in GLU and 7,366 sites in OLIG. Fourth, we confirmed a common local sequence motif for all known and novel A-to-G editing events, whereby guanosine is depleted - 1bp upstream and enriched +1 downstream the targeted adenosine, as previously reported (**Figure 2F-G**). Notably, novel A-to-G sites displayed significantly more supporting read coverage compared to known sites (**Figure 2H**) and exhibited consistent editing rates of ∼37% on average across all cells (∼3% less than known sites, *p*<2.16×10^−20^, linear regression) (**Figure 2I**). Moreover, 7,326 novel A-to-G sites validated across two or more cell types (**Figure 2J**). We also identified more than 20,000 novel editing sites with substitution types other than A-to-G. Of these, ∼42% were C-to-T and G-to-A edits, which we treated as provisional (**Supplemental Table 2**).

### Partitioning the variance in RNA editing rates explained by known factors

We studied 15,221 A-to-G sites detected across all three cell types and all donors to quantify the fraction of RNA editing variance explained by eight known biological and technical factors. Collectively, these factors explained ∼28% of RNA editing variation. Differences between cell types had the largest genome-wide effect, explaining a median ∼8.3% of the observed variation, followed by differences in chronological age (∼6.7%), pH (∼3.5%), *ADAR2* (∼1.8%) and *ADAR1* expression (∼1.5%) (**Figure S3A**). Donor as a repeated measure had a small but detectable effect for ∼6% of editing sites. As expected, principal component analysis (PCA) accurately distinguished MGE-GABA and GLU neurons from OLIG along the first PC1, explaining 29.2% of the variance (**Figure S3B**). Using a linear regression model, we also catalogued a total of 6,765, 7,703 and 2,540 RNA editing sites that were significantly associated with *ADAR1*, *ADAR2* or *ADAR3* expression after adjusting for repeated measures (FDR < 0.05), respectively (**Figure S3C, Supplemental Table 3**).

In addition to ADAR, several RNA binding proteins (RBPs) can also act as global mediators of RNA editing. We found a total of 470 RBPs were differentially expressed (FDR < 0.05, log fold-change > 0.5), and roughly half (∼49%) were more highly expressed in OLIG relative to GLU and MGE-GABA (**Figure S4A-B**). A total of 170 OLIG-specific RBPs, including *NOP14*, *PTEN*, and *DYNC1H1* were negatively correlated with global editing activity, while 161 neuron-specific RBPs, including *MOV10*, *CELF4*, and *FMRP* had a positive association with global editing rates (**Figure S4C, Supplemental Table 3**). We further examined whether FMRP binding sites were enriched near MGE-GABA, GLU and OLIG editing sites using enhanced ultraviolet crosslinking and immunoprecipitation (eCLIP) in human frontal cortex. *FMRP* eCLIP peaks were significantly enriched for MGE-GABA (permutation *p*-value=0.005, Z-score=9.3) and GLU sites (permutation *p*-value=0.005, Z-score=6.2), and moderately enriched for OLIG sites (permutation *p*-value=0.005, Z-score=7.4) (**Figure S4D-E**). Thus, these RBPs may work alongside ADARs as *trans* regulators of editing levels in brain cell populations.

### Cell type-enriched RNA editing sites

To identify quantitative differences in RNA editing levels among sites detected across two or more cell types, we computed three pairwise comparisons (*i.e.,* MGE-GABA vs. GLU, MGE-GABA vs. OLIG, GLU vs. OLIG) and adjusted each linear model for the possible influence of post-mortem interval (PMI), age, and donor as a repeated measure (**Figure 3A, Supplemental Table 4**). Overall, we identified 9,022 “cell type-enriched” sites which displayed significantly higher editing in one cell population relative to another. The majority of these sites were more highly edited in MGE-GABA and GLU relative to OLIG (**Figure 3B**), are previously reported editing sites, and commonly mapped to introns, 3’UTRs and intergenic regions (**Figure 3C**). Notably, after adjusting for *ADAR1* and *ADAR2* expression, we found substantially fewer differentially edited sites (*n*=192 sites), suggesting the differential editing at the majority of these sites depends on ADAR activity (**Supplemental Table 4**). Functional annotation revealed that cell type-enriched sites in each cell type were associated with genes involved in neuronal differentiation and cell adhesion, but were also enriched for unique processes (**Supplemental Table 4**). MGE-GABA sites were uniquely enriched for genes implicated in trans-synaptic signalling, MAPK signaling, as well as genes at the postsynaptic density (**Figure S5**). GLU sites were uniquely enriched for genes implicated in chromatin organization and interferon signaling. OLIG sites were uniquely enriched for genes associated with N-acetyltransferase activity and methylated histone binding. Notably, cell type-enrichment differences in RNA editing rates were not strongly associated with cell-specific differential gene expression (**Figure S6**).

**Figure 3.**
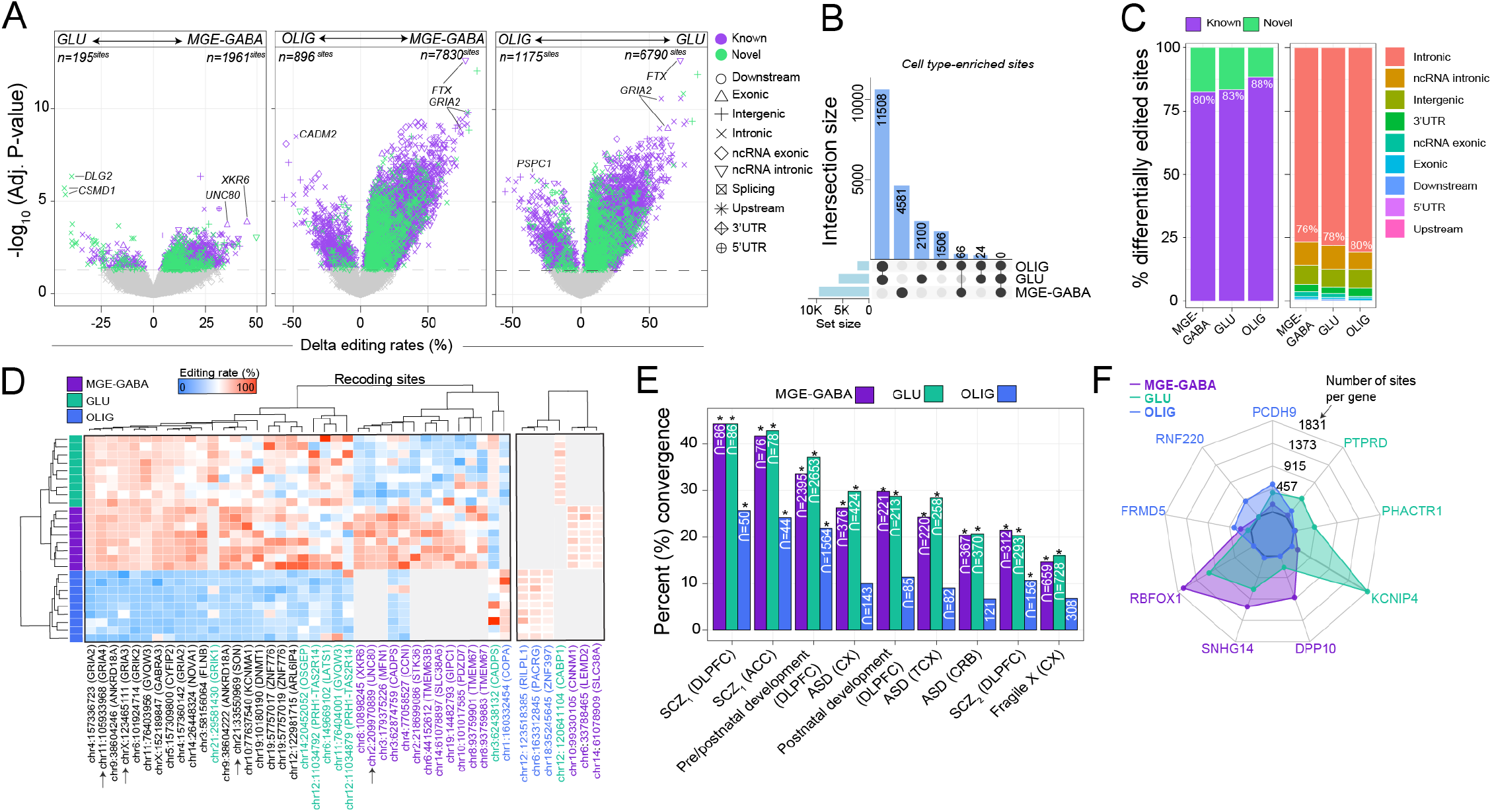
Quantifying variance in RNA editing rates. **(A)** Differential RNA editing analysis was computed in a pairwise fashion between all three FANS-derived cell populations, including GLU vs. GABA (left), OLIG vs. MGE-GABA (center) and OLIG vs. GLU (right). RNA editing sites are uniquely shaped by genic region and colored by novelty. **(B)** UpSet plot reporting counts for all cell type-enriched sites are reported with their overlaps. **(C)** The percentage (%, y-axis) of differentially edited sites by novelty (left) and genic region (right). **(D)** Heatmap displaying differential editing levels at recoding sites across MGE-GABA (blue), GLU (purple), and OLIG (green) cells. Grey color indicates sites that were not detected in a given cell type. Arrows pointing to specific sites indicate sites validated by independent methods (see Figure S8). (**E**) The convergence (%, y-axis) for each cell-type specific set of editing sites was compared to several published reports of editing sites dysregulated in bulk brain tissue (*see Methods*). A Fisher’s exact test and an estimated odds-ratio was used to compute significance of the overlap. (**F**) Spider plot of the top three genes for each cell type that display enrichment for A-to-G RNA editing sites after accounting for gene length.

A small fraction of differentially edited sites was catalogued as RNA recoding events (∼0.4%, *n*=38 sites), which alter amino acid states and displayed increased conservation relative to sites in other genic regions (**Figure S7, Supplemental Table 5**). These recoding events were mainly more highly edited in neurons relative to OLIG and included several well-known sites involved in tight regulation of Ca^2+^ permeability (Q➝R *in GRIA2*,), actin cytoskeletal remodeling at excitatory synapses (K➝E *in CYFIP2*) and gating kinetics of inhibitory receptors (I➝M *in GABRA3*) (**Figure 3D**). Twelve recoding sites were more frequently edited in MGE-GABA compared to GLU, including an R➝G site in cyclin-I (*CCNI*), a K➝R site in mitofusin 1 (*MFN1*), a E➝G site in calcium dependent secretion activator (*CADPS*). Notably, one I➝V site in coatomer subunit alpha (*COPA*) was more highly edited in OLIG relative to neurons. We tested the cellular specificity for four recoding sites using an independent method (site-specific PCR amplification of regions harboring editing sites followed by sequencing) applied to orbitofrontal cortex samples from six independent adult donors (**Figure S8, Supplemental Table 6**). We confirmed three R➝G sites in SON DNA binding protein (*SON*), *GRIA3* and *GRIA4*, which were more highly edited in neurons relative to OLIG. We also validated one S➝G site in *UNC80,* a component of the NALCN sodium channel complex (*UNC80*), which was more highly edited in MGE-GABA neurons.

We explored whether our data can resolve the cellular specificity of editing sites previously described to be dysregulated in bulk brain tissue across neurodevelopment and in neurological disorders (**Figure 3E, Supplemental Table 4**). We observed a strong enrichment of MGE-GABA and GLU sites among disease-linked sites, most notably in dorsolateral prefrontal cortex (DLPFC) and anterior cingulate cortex (ACC) of schizophrenia patients (*p*=0.009, *p*=6.3×10^−5^, respectively). Also, MGE-GABA and GLU sites were enriched for sites in the DLPFC previously found to be dynamically regulated throughout prenatal and postnatal development (*p*=6.1×10^−279^, *p*=2.3×10^−214^, respectively). Enrichment for OLIG RNA editing sites was consistently lower across all independent studies and cohorts, as expected. These results reconfirm the validity of our approach and shed light onto some of the cellular origins of altered RNA editing in neurodevelopment and disease^17–22^.

### Genes enriched for RNA editing sites within cellular populations

We found an association between gene length and the number of RNA editing sites per gene within MGE-GABA (*R^2^*=0.16), GLU (*R^2^*=0.15) and OLIG cells (*R^2^*=0.04) (**Figure S9A, Supplemental Table 7**). After normalizing the total number of RNA editing sites by gene length (*see Methods*), we found a higher density of RNA editing sites in OLIG cells at several genes associated with OLIG-specific expression: *PCDH9* (*n*=560 sites), *RNF220* (*n*=419 sites), *FRMD5* (*n*=277 sites) (**Figure 3F**, **Figure S9B-C**). In MGE-GABA, genes *RBFOX1* (*n*=1696 sites), *SNHG14* (*n*=1083 sites) and *DPP10* (*n*=243 sites) were enriched for editing sites. In GLU, genes *KCNIP4* (*n*=1831 sites), *PHACTR1* (*n*=433 sites) and *PTPRD* (n=488) were enriched (**Figure 3F, Figure S9B-C, Supplemental Table 7**). Notably, genes with higher rates of editing in MGE-GABA and GLU cells were not associated with cell-specific differential gene expression.

### Validation of RNA editing sites by independent snRNA-seq data

We next examined RNA editing across all brain cell types using single-nuclei RNA sequencing (snRNA-seq) of the adult PFC generated by PsychENCODE^30^ (*n*=3 biological replicates). A total of 24 discrete cell clusters comprising six major cell types were identified through unsupervised dimension reduction and annotated using previously defined cell marker genes (**Figure 4A**). To overcome data sparsity associated with snRNA-seq data, we binned nuclei into pseudo-bulk pools that most closely reflect the MGE-GABA, GLU and OLIG populations in our FANS datasets based on the expression of their markers *SOX6*, *RBFOX3*, and/or *SOX10,* respectively (**Figure 4B, Figure S10**). We observed that ∼52% of all nuclei expressed the GLU marker (*n*^nuclei^=8957), ∼21% expressed the MGE-GABA marker (*n*^nuclei^=3708), and ∼9% expressed the OLIG marker (*n*^nuclei^=1709). Notably, a small subset of which were assigned to the GLU pseudo-bulk pool were also positive for markers of CGE-derived inhibitory neurons (e.g. *VIP* and *LAMP5*), which make up ∼7% of nuclei overall. This was consistent with our FANS strategy, which separated MGE-GABA form other neurons using the MGE-GABA specific marker SOX6. Thus, a small population of non MGE-derived GABA neurons (∼10-12% of all GABA neurons) was sorted together with GLU neurons^29^. We generated three additional pseudo-bulk pools representing astrocytes (*n*^nuclei^=2,078), endothelial cells (*n*^nuclei^=464) and microglia (*n*^nuclei^=130). We examined ADAR expression and global editing rates within each the six cellular pool (*see Methods*), confirming higher global editing activity as well as higher expression of *ADAR1* and *ADAR2* in MGE-GABA and GLU relative to OLIG cells (*p*=0.002, linear regression), and relative to all remaining non-neuronal cell types (*p*=0.0002) (**Figure 4C**).

**Figure 4.**
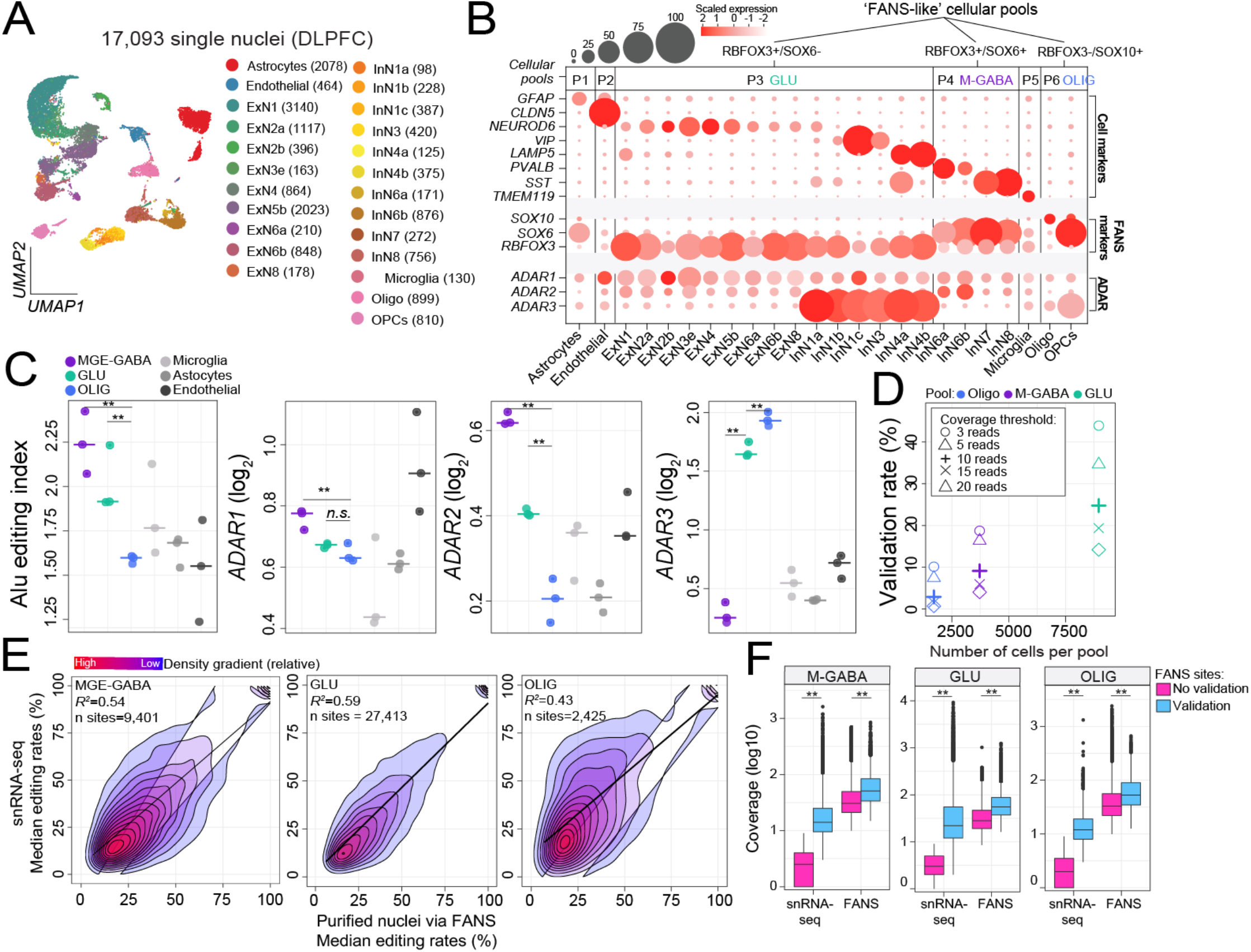
Confirmation of cell-specific editing in snRNA-seq. **(A)** snRNA-seq performed on adult PFC (*n*= 3) classified 24 unique cell populations. Values in brackets indicate number of cells per subset. **(B)** Cells were further collapsed into six major cell classes, which were evaluated for expression of our FANS-derived cell populations (*RBFOX3*, *SOX6*, and/or *SOX10*) to identify cells that define ‘FANS-like cellular pools’. **(C)** The AEI and expression profiles for *ADAR1*, *ADAR2*, and *ADAR3* (y-axes) were computed for each ‘FANS-like cellular pool’ in addition to microglia, astrocytes, and endothelial cells. **(D)** The percentage of RNA editing sites (y-axis) from the original FANS-derived cell populations that validated in each respective ‘FANS-like pool’ was determined across varying read coverage thresholds. **(E)** Comparison of the median editing rates (%) within each cell-type for sites validated by both approaches with a minimum coverage of 10 reads illustrates correlation of the two approaches. **(F)** Sites which failed to validate by snRNA-seq were largely explained by difference in read coverage and total number of cells sequenced.

Furthermore, we queried all selective RNA editing sites derived by FANS (**Figure 2**) within each snRNA-seq cellular pool and validated 9,401 sites in MGE-GABA (∼8%), 21,413 sites in GLU (∼27%) and 2,425 sites in OLIG (∼3%) pools with high-confidence (**Supplemental Table 8**). Of these sites, a total of 1,748, 4,315 and 231 were deemed novel sites in MGE-GABA, GLU and OLIG populations, respectively. Notably, validation rates depended on snRNA-seq coverage thresholds and the number of nuclei included within each pseudo-bulk pool, with larger pools having higher validation rates (*e.g.* GLU) (**Figure 4D**). Nevertheless, for the sites passing the defined coverage thresholds, we observed a high level of concordance between editing rates quantified in purified nuclei via FANS relative to editing rates quantified via snRNA-seq for MGE-GABA (*R^2^*=0.54), GLU (*R^2^*=0.59) and OLIG (*R^2^*=0.43) (**Figure 4E**). RNA editing sites that did not validate in snRNA-seq were largely explained by the lack of supporting read snRNA-seq coverage or lower expression in FANS (**Figure 4F**).

### Imputing cell-type specific RNA editing rates in bulk RNA-seq

The challenges of cell type purification and single cell sequencing have precluded broad application of those techniques. To leverage the extensive bulk RNA-seq datasets generated from human brains, we sought to use our data to infer cell type specific RNA editing in 1,129 bulk RNA-seq samples across 13 brain regions from the Genotype-Tissue Expression (GTEx) project (**Supplemental Table 9**). Given that global editing activity is highest in neurons, we anticipated that a substantial fraction of the variation in global selective A-to-G editing rates in bulk brain tissue would be explained by the proportions of neurons within each sample. Moreover, given the high proportion of neurons in the cerebellum, we expected that editing rates would be highest in this region relative to all others. We used cell type deconvolution to estimate the proportions of six major CNS cell types for each bulk RNA-seq sample (*see Methods*).

We observed considerable brain region variability in global selective editing activity, defined by the AEI. The cerebellum and cerebellar hemisphere had higher editing activity than all other brain regions (*p*=2.7×10^−1^^10^, Cohen’s d=3.00) (**Figure 5A**). Similarly, cellular deconvolution of all bulk tissue samples confirmed elevated neuronal proportions in cerebellum and cerebellar hemisphere relative to all other regions (*p*=1.3×10^−1^^32^, Cohen’s d=5.65). Importantly, a significant fraction of the variance in global editing activity was explained by differences in neuronal fractions both within and across all brain regions (*R^2^*=0.31; **Figure 5B**). Donors and regions with higher proportions of neurons displayed elevated global editing activity. Notably, other biological and technical factors were unable to explain as much variation (**Figure S11**). Using these bulk RNA-seq data, we also confirmed the positive association between the proportion of neurons and *ADAR1* (*R^2^*=0.55) and *ADAR2* expression (*R^2^*=0.70), and a negative association with *ADAR3* (*R^2^*=-0.22) (**Figure 5C**). For further context, we performed PCA on *ADAR1*, *ADAR2* and the AEI metric, which accurately stratified all samples by brain region and the proportion of neurons (**Figure S12**). Collectively, these results confirm the cellular specificity of ADARs and the AEI in bulk tissue as observed in our FANS-derived nuclei.

**Figure 5.**
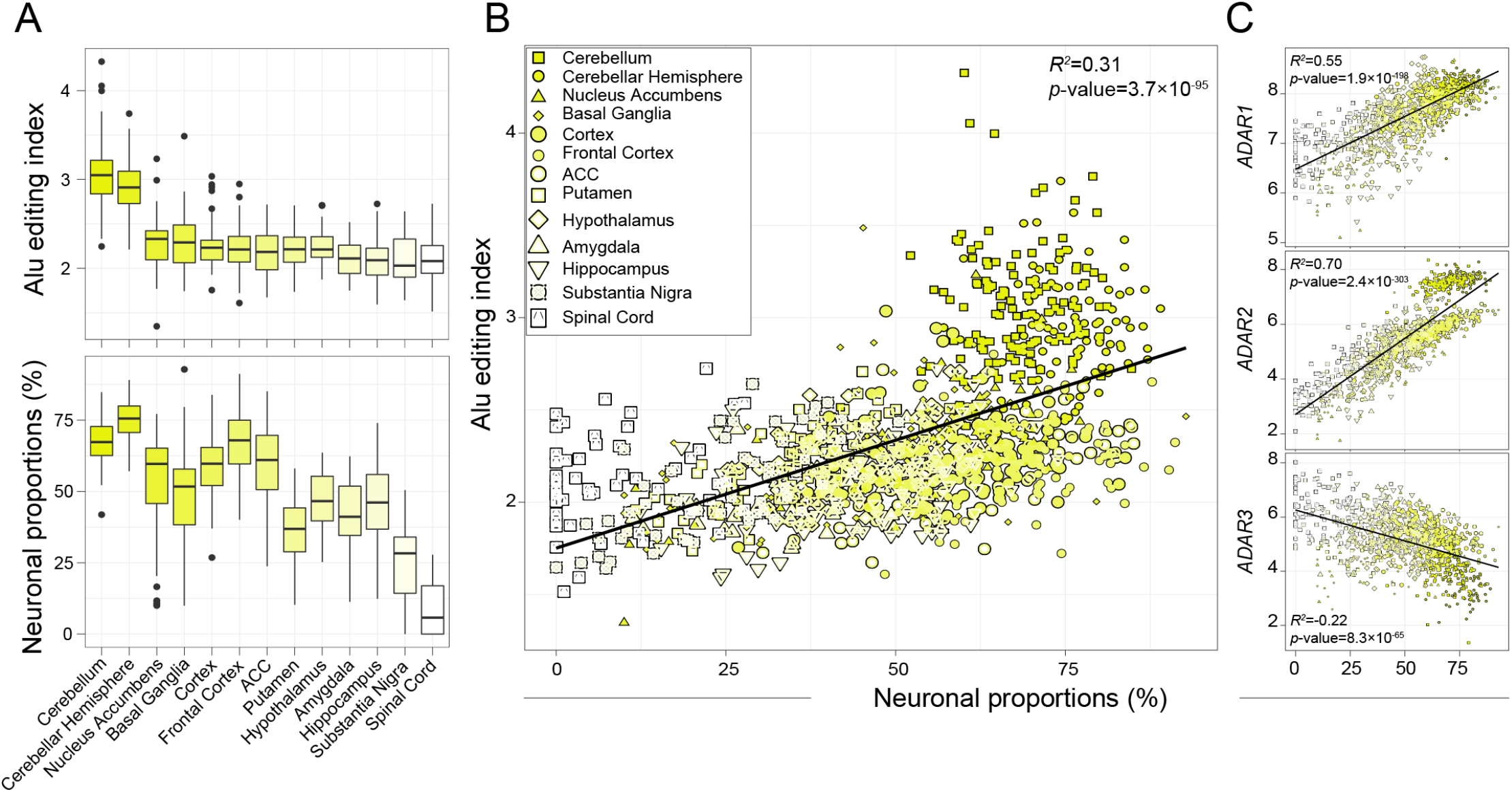
Variance in global editing activity in bulk tissue explained by neuronal content. **(A)** Alu editing index (AEI) (y-axis, upper) was computed across 13 brain regions from the GTEx consortium projects and neuronal cell type proportions (y-axis, lower) were estimated. Cellular deconvolution was applied using the Darmanis et al., 2015 reference signature matrix. **(B)** The amount of AEI variance and (**C**) ADAR expression explained by neuronal proportions was determined using a linear regression model across all 13 brain tissues.

Next, we queried RNA editing sites in all GTEx bulk brain samples based on a list of known sites including our 189,229 bona fide cell type-associated sites plus all other sites listed in the REDIportal database. As expected, the number of editing sites in the cerebellum and cerebellar hemisphere (∼58,143 per donor) was threefold greater than all other regions (∼18,456 sites per donor, *p*=1.3×10^−1^^32^, Cohen’s d=2.99) (**Figure 6A**). Intra-donor variation in site detection was similarly associated with the proportion of neurons (*R^2^*=0.70), *ADAR2* (*R^2^*=0.48) and *ADAR1* (*R^2^*=0.30) expression (**Figure S13**). RNA editing sites mapped primarily to 3’UTRs (∼40%) and introns (∼22%) with few recoding events (∼2.9%) across regions (**Figure S14A**). Notably, ∼20% of all detected sites per donor were catalogued as either cell type-specific or -enriched in either MGE-GABA, GLU and/or OLIG cells and these sites displayed significantly higher detection rates in the cortical regions relative to all other regions (*p*=2.8×10^−28^, linear regression) (**Figure 6B**). Importantly, these sites also had twentyfold higher editing rates relative to detected sites that were catalogued in the REDIportal database but not by FANS (**Figure 6C**). Moreover, novel sites identified by FANS were detected at an expected low rate in bulk tissue (∼213 novel sites on average across all donors and GTEx regions) (**Figure S14B**).

**Figure 6.**
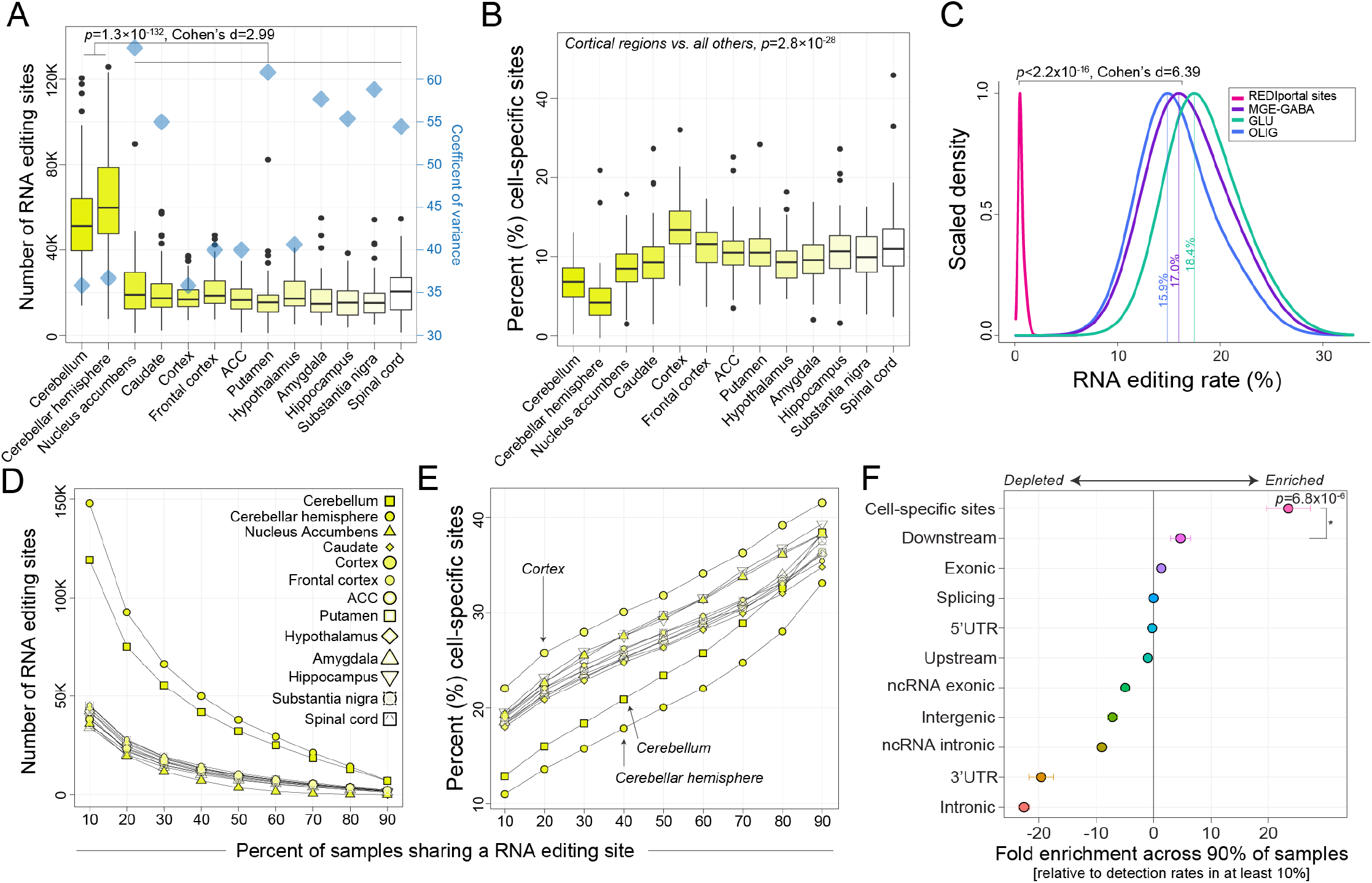
Annotation of cell type-associated sites in bulk tissue. **(A)** Total number of editing sites (y-axis) identified is reported along with coefficient of variance (blue diamond) for each GTEx brain tissue to describe intraregional fluctuation observed amongst donors. **(B)** The percentage of cell type-associated RNA editing sites out of total number of editing sites per donor (y-axis), with a clear enrichment in the cortex. **(C)** Density distribution plots delineate that the editing levels of cell type-associated relative to all other sites detected in bulk tissue. **(D)** A sliding threshold was used to identify sites shared amongst a given percentage of samples per region (x-axis). **(E)** Following each threshold, the fraction of cell type-associated RNA editing sites were computed, further highlighting cortical enrichment. (**F**) Fold-enrichment analysis informs cell type-associated sites are most commonly detected across the majority of donors.

Given the substantial inter-donor variability in the number of detected RNA editing sites (**Figure 6A, Figure S13**), we asked whether cell type-specific and -enriched sites were consistently detected and edited across the majority of samples (**Figure 6D**). The number of commonly edited sites substantially decreases when gradually increasing the requirement of a site to be detected in larger fraction of donors per region in an incremental fashion (**Figure 6D**). Notably, following each iteration, the proportion of retained sites that identify as cell type-associated RNA editing sites gradually increased (**Figure 6E**), indicating that such sites are detectable within bulk RNA-seq tissues across the vast majority of donors. We tested this result by computing fold enrichment for all cell type-associated sites relative to sites assigned to specific genic regions and found that these editing sites were disproportionally enriched across the majority of samples (*p*=6.8×10^−6^), followed by RNA editing sites mapping to downstream transcription start site positions, sites in exons and those in splice-site regions (**Figure 6F, Supplemental Table 10**). These results suggest that sites with cellular resolution comprise those that are most commonly detected across the majority of donors and regions in bulk tissue.

### Genetic variants affect the rate of selective RNA editing

We used imputed genotype data from the GTEx project to detect common single nucleotide polymorphisms (SNPs) that associate with RNA editing levels (edQTL, editing quantitative trait loci) (*see Methods*). To identify genetic variants that could explain the variability of selective RNA editing, we ran association tests across each brain region and identified 661,791 cis-edQTLs (*i.e.* SNPs located within 1 Mbp of an RNA editing site) at a genome-wide FDR < 5% (**Figure 7A**). Each max-edQTL (defined as the most significant SNP-site pair per site, if any) meeting a genome-wide significance (FDR < 0.05) was located close to their associated editing site and acting in cis (200kb ± bp the target adenosine) (**Figure 7B**). Max-edQTLs were examined for cellular specificity according to the aforementioned analyses. A total of 5,011, 4,514, and 3,677 cis-edQTLs were annotated as either MGE-GABA, GLU and/or OLIG associated, respectively. We found hundreds of cis-edQTLs in each brain region, with an especially large number in the cerebellar regions (**Figure 7C**). Overall, a total of 13,438 unique editing sites (eSites) displayed edQTLs and these sites were predominately located in 3’UTRs (∼38%) (**Figure 7C**). Of these, 1,869 eSites (∼13.9%) were genetically regulated across three or more brain regions, while 51 eSites corresponding to 47 unique genes were commonly regulated across all thirteen regions (**Figure 7D, Supplemental Table 11**). For example, an RNA editing site located on the 3’UTR of glutathione-disulfide reductase (*GSR*), a site which shows strong preferential editing in neurons, is consistently associated with the same SNP (chr8:30536581) across all thirteen brain regions with similar direction of effect (**Figure 7E**). Importantly, our eSites enriched for 618 out of 977 eSites previously identified across GTEx brain regions^27^ (*p*=2.1×10^−31^, Fisher’s exact test) and expand upon these efforts by 13-fold (**Supplemental Table 11**).

**Figure 7.**
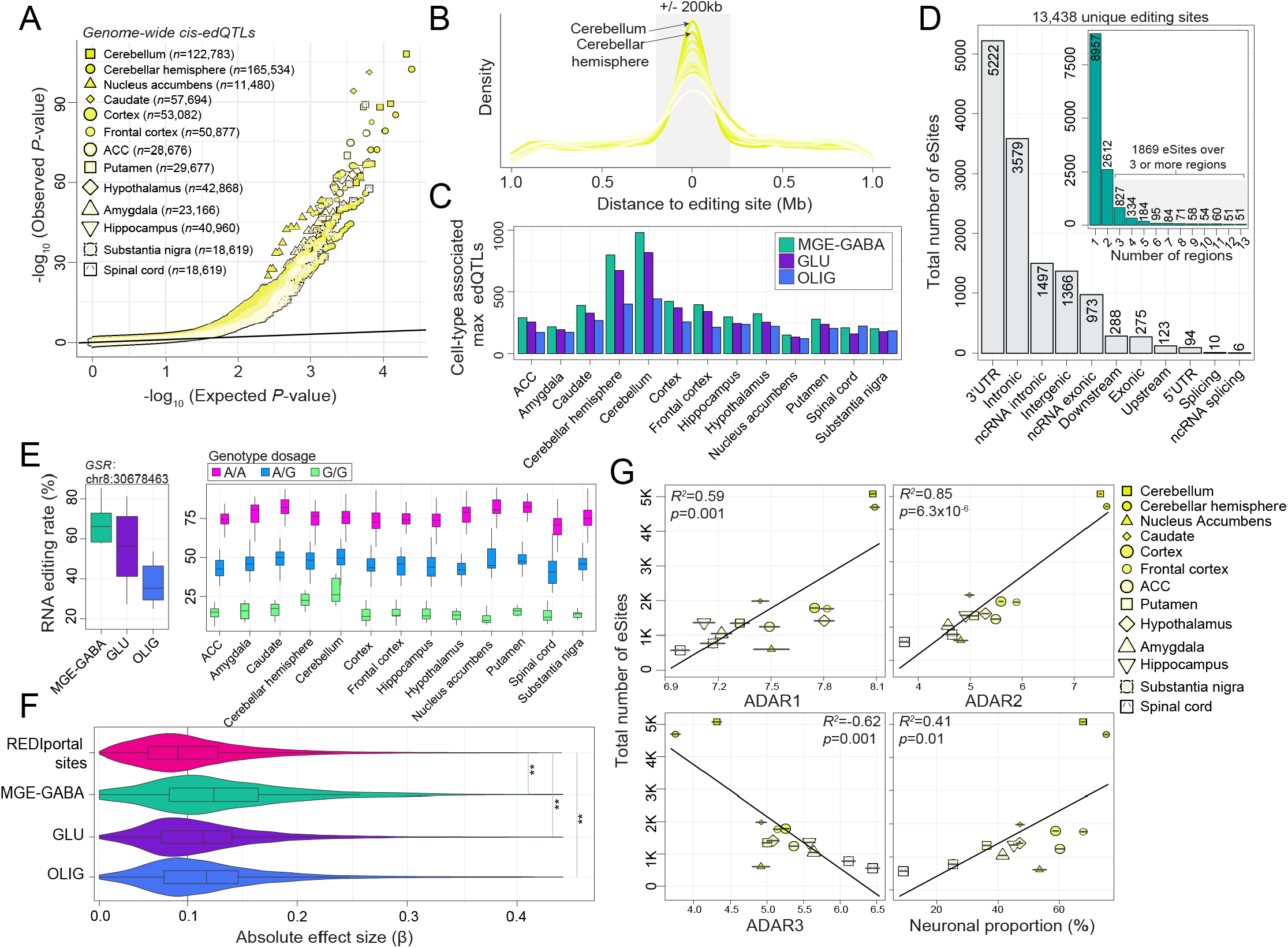
Identification and annotation of edQTLs across brain regions. **(A)** QQ-plot depiction of observed versus expected *P*-values for genome wide cis-edQTLs**. (B)** Max edQTLs are highly enriched within +/-200bp of their respective editing sites. **(C)** Counts of cell type-associated max edQTLs in each brain region. (**D**) The total number of eSites (unique editing sites with associated edQTL) are broken down by genic region. Inset plot counts the number of eSites commonly detected across more than one region. (**E**) An example of one commonly detected edQTL across all regions with neuronal cell type enriched editing rates (left) with consistent direction of edQTL effect across all regions (right). (**F**) The absolute effect size (β, x-axis) is compared for edQTLs annotated as cell type-associated or edQTLs without cellular identify. (**G)** Regression analysis indicates that eSite discovery is dependent upon regions expressing higher levels of ADAR1, ADAR2 and those with higher proportions of neurons.

We also compared the absolute effect sizes across all max-edQTLs and observed significantly stronger associations for cell type-specific max-edQTLs relative to those with RNA editing sites that were detected in REDIportal but not by FANS (**Figure 7F**). These results were consistent across brain regions (**Figure S15**). Notably, max-edQTL SNPs were moderately enriched in brain-specific enhancers and promoters across regions (**Supplemental Table 11**). Additionally, while sample size is the gold-standard metric to determine power and discovery for QTL studies (**Figure S16**), the number of unique eSites detected per brain region also correlated with *ADAR1* (*R^2^*=0.59) and *ADAR2* (*R^2^*=0.85) expression, as well as with the proportion of neurons per brain region (*R^2^*=0.41) (**Figure 7G**). We confirm these associations using a smaller set of results from an independent analysis of eSite discovery across GTEx brain regions^27^ (**Figure S17**). These results suggest that, in addition to analyzing large sample numbers, edQTL discovery is highly context dependent: tissues with higher levels of ADAR expression and increased neuronal content provide favorable frameworks for discovery.

## DISCUSSION

Exposing highly regulated editing sites across different cell populations, brain regions and those that are genetically regulated can promote studies to dissect their functional relevance within the right cellular context. Given the dearth of studies examining cellular features of RNA editing in the brain, we first set out to increase the resolution of RNA editing among three main cell populations in the human cortex. Through FANS, unique populations of cell types were isolated from the cortex with distinct functional differences. Using RNA-seq, we determined how RNA editing facilitates transcriptomic diversity more commonly in MGE-GABA and GLU neuronal populations relative to the major glial population in the brain – OLIG cells, documenting numerous sites and genes with cell-specific editing properties. snRNA-seq data was used to validate global trends in editing among CNS cell types and confirm the cellular resolution for a subset of RNA editing sites and genes. Subsequently, we applied rules of cell-specific RNA editing to quantify variability in RNA editing rates observed in bulk brain tissues from the GTEx project, and successfully show that: *i)* tissues and donors with higher neuronal content display increased global editing rates and exhibit increased detection rates of RNA editing sites; *ii)* commonly detected sites quantified in bulk brain tissue can be classified as cell type-specific or -enriched RNA editing sites; *iii)* hundreds of edQTLs classify as cell type-specific or -enriched and display larger effect sizes than edQTLs without cellular resolution; and *iv)* similar to RNA editing site detection, edQTL detection is also better powered in regions that express higher levels of *ADAR2* and display increased neuronal content. These results illuminate new differences in RNA editing sites and rates among cortical cell populations and establish new frameworks for future large-scale studies of bulk RNA-seq brain tissue to interpret RNA editing sites and their variability.

We observed increased global editing rates in neurons relative to OLIG populations, similar to previous reports among CNS cell types in mice and *Drosophila*^23–25^. These global trends were accompanied by increased detection in the number of RNA edited sites in neurons. Such differences were largely associated with differences in ADAR expression levels, rather than being driven by differential expression of the edited transcripts. We also uncovered a several editing sites in close proximity on the same transcript and co-regulated in the same cell population (*e.g.* editing sites on *CSMD1* in OLIG, *KCNIP4* in GLU and *RBFOX1* in MGE-GABA), groups of sites which illustrate a regulation of editing that can exert its effect differently in different parts of the same transcript, as recently shown across neuronal populations in *Drosophila*^25^. While such events would be successively edited by ADAR, regulation by RBPs may also have a similar effect^31^. Several RBPs were preferentially expressed in either MGE-GABA and GLU populations or OLIG and were either positively or negatively correlated with global editing activity. Such RBPs might explain a fraction of the differential RNA editing levels observed at different sites in distinct populations. However, follow-up functional validation is required to fully understand the *trans* regulation of editing levels by RBPs among brain cell types.

Our study not only provides cellular context for already existing sites in the brain, but also identified several previously unknown editing sites likely masked by bulk RNA-seq sampling techniques. For example, a known recoding site (S➝G) in *UNC80* was preferentially edited in MGE-GABA neurons (**Figure 3D, Figure S8**). This gene is essential for NALCN sensitivity to extracellular Ca^2+^ and mutations in *UNC80* are associated with congenital infantile encephalopathy, intellectual disability and growth issues^32^. Moreover, novel sites, which were often found in moderate-to-lowly expressed genes, mostly overlapped Alu elements of the transcriptome, consistent with existing reports^3, 4, 32, 33^. Further, our nuclei data, perhaps unsurprisingly, indicate that ADAR driven RNA editing activity occurs predominately in non-coding transcripts, also consistent with existing reports^3, 4, 33^. Such editing sites may regulate circular RNA biogenesis^34^ or RNA interference pathways^35^, which in turn can alter heterochromatin formation. While follow-up investigations are required to elucidate the functional relevance of these editing sites, these data indicate that transcriptome sequencing of cell populations purified by FANS can accelerate the discovery of novel sites, including difficult to detect transcripts.

snRNA-sequencing data was used to confirm global RNA editing trends among MGE-GABA, GLU and OLIG populations, and shed additional light on global editing patterns across three additional CNS cell types. We also validated several thousand cell-specific RNA editing sites using an *in silico* cellular pooling technique to overcome hurdles related to snRNA-seq. Specifically, calling RNA editing sites from snRNA-seq is challenging as the technique is limited from extremely low capture efficiency and low sequencing depth with reads covering only a fraction of the entire genome^36, 37^. Moreover, intronic regions, which were highly edited in FANS-derived data, often represent high vulnerability to low-coverage in scRNA-seq experiments, where detection of A-to-I events is reduced^38^. Validation rates by snRNA-seq were highest for GLU and MGE-GABA populations, which constituted pools with the largest number of nuclei, and concordance of editing levels for these sites was exceptionally high between snRNA-seq and FANS-derived data sets. Until now, there has been limited investigation of the accuracy of RNA editing calls from snRNA-seq data^26^. We anticipate that future work applying statistical models to snRNA-seq to quantify editing will benefit from increased sample sizes (*i.e.* more biological replicates) and longer, more deeply sequenced reads, which may also award the detection of novel sites necessary to build cell-specific profiles.

Our results also underscore several important features of RNA editing in bulk brain RNA-seq, and in doing so, highlight the tremendous heterogeneity of RNA editing site quantification and detection at population scale. The majority of randomly selected sites are detected only across a subset of donors within a particular brain region, and these detection rates were largely a function of *ADAR2* expression and whether a region exhibited increased neuronal content (similar to variation in the AEI in bulk tissue). Differences in pools of cellular RNA across samples and randomly sequenced RNA fragments offer likely explanations to such intra-donor variability in site detection. Another possible explanation to account for such heterogeneity is that RNA editing is a transient process and such differences may simply reflect distinctions in the timing of cellular processes and signaling cascades. Indeed, at the level of individual cells, dynamic responses to environmental cues occur on a timescale that is faster than activity-induced transcription via the coordinated, activity-induced switching of internal molecular states and cellular metabolism^1, 2^. Moreover, we detected 13,438 unique eSites, which occurred predominantly in 3’UTRs across brain regions. While this level of discovery significantly expands upon existing counts of genetically regulated editing sites in the brain^27^, we anticipate that this number is still an underestimate. Our edQTLs do not include rare RNA editing sites that were detected in a small subset of samples (which are the majority) that may also be genetically regulated. Future work considering edQTL models that account for rare editing sites may lead to a more resolved atlas of RNA editing sites and their genetic regulation in the brain.

Finally, while our data provide new avenues for understanding cellular specificity RNA editing, one key question remains: ‘*What is the biological explanation as to why neurons exhibit a preponderance of RNA editing activity?*’. Here we extend some putative solutions. One possible explanation of enhanced RNA sequence diversity and ADAR expression in neurons might be to afford increased neural plasticity. Neurons, more than other cell types, must be able to quickly respond to altered environmental inputs, therefore, RNA editing may represent a mechanism capable of aligning with the timing required for experience-dependent plasticity, similar to other RNA modifications^39, 40^. As such, editing activity in neurons might play a role in controlling the organizational architecture of neuronal networks implicated in higher order learning and memory. A related explanation of enhanced diversity of RNA editing in neurons may hint at its potential relation with essential neuronal functions.

This idea is reinforced by the high density of editing in neuronal transcripts that encode proteins directly involved in excitatory/inhibitory (E/I) functions and the significant differences in editing between excitatory GLU and inhibitory GABA neurons. Such differences may modulate E/I balance within the cortical circuitry. Imbalance of E/I activity is thought to play a critical role in the pathophysiology of several different neuropsychiatric and neurological disorders, including autism spectrum disorder, schizophrenia, and epilepsy^41, 42^. Thus, functional relevance of the observed RNA editing differences between different populations of brain cells in health and disease warrants further investigation. Lastly, it remains unclear how RNA editing may play out across distinct cellular compartments and locations. Recent work has found discrete gene expression differences across synapses, dendrites, axons and neuronal bodies^43, 44^, and RNA editing may represent a driver of activity-induced RNA localization^1, 2, 11^.

## MATERIALS AND METHODS

### Experimental design and RNA-sequencing samples

The following RNA-seq datasets were leveraged to quantify RNA editing. Raw FASTQ files or mapped bam files of the following datasets were downloaded from either the sequencing read archive or Synapse.org:

1. **FANS-derived cortical cell populations**: Raw FASTQ files were obtained for a total of 27 paired-end (125 bp) nuclei samples were collected from MGE-GABA (*n*=9), GLU (*n*=9) and OLIG (*n*=9) populations (syn12034263). Antibodies against brain cell population markers NeuN, SOX6, and SOX10 were used in the FANS protocol, as described in our previous publication^29^. Briefly, NeuN (also known as RNA-binding protein *RBFOX3*) is a well-established marker of neuronal nuclei; *SOX6* is a transcription factor expressed in MGE-GABA neurons during development and into adulthood; the use of anti-SOX6 antibodies to separate the nuclei of MGE-GABA from GLU neurons was described by us previously^45, 46^; *SOX10* is a transcription factor specifically expressed in OLIG.
2. **Single-nuclei RNA-sequencing**: Mapped bam files were obtained for snRNA-seq generated from 17,093 nuclei from the dorsolateral prefrontal cortex (DLPFC) of three adult brains (syn15672826). The 10X Genomics chromium platform was used to capture and barcode single nuclei using the Chromium Single Cell 3’ Library and Gel Bead Kit v2 (10x Genomics) and the Chromium Single Cell A Chip Kit (10x Genomics) as previously described^30^.
3. **Transcriptome and genotype data from Genotype-Tissue Expression Project:** We obtained approval to access Genotype-Tissue Expression (GTEx) Project through the database of Genotypes and Phenotypes (dbGaP) (phs000424.v8). Raw FASTQ files were obtained for a total of 1431 paired-end (75 bp) samples were collected across 13 brain regions (anterior cingulate cortex (ACC), amygdala, caudate, cerebellar hemisphere, cerebellum, cortex, frontal cortex, hippocampus, nucleus accumbens, putamen, spinal cord, substantia nigra). The VCFs for the imputed array data were available through dgGAP, in phg000520.v2.GTEx.MidPoint.Imputation.genotype-calls-vcf.c1.GRU.tar (the archive contains a VCF for chromosomes 1-22 and a VCF for chromosome X).

### Identification of site-selective RNA editing sites

All FASTQ files were mapped to human reference genome (GRCh38) using STAR v2.7.3^47^ and mapped files were used as input for the subsequent analysis. To quantify site-selective RNA editing sites from FANS-derived cortical cell populations, we used a combination of *de novo* calling via Reditools v2.0^48^ (parameters: -S -s 2 -ss 5 -mrl 50 -q 10 -bq 20 -C -T 2 –os 5) and supervised calling of known RNA editing sites from the REDIportal database^49^ as a second pass. A number of filtering steps were applied to retain only high-quality, high-confident bona fide RNA editing sites: i) all multi-allelic events were discarded; ii) a minimum total read coverage of 10 reads and at least 3 edited reads was required to classify as an editing event; iii) any sites mapping to homopolymeric regions or in hg38 blacklisted regions of the genome^50^ were discarded; iv) any sites mapping to common genomic variation in dbSNP(v150) and those in gnomAD with minor allele frequency greater than 0.05 were discarded; v) for FANS-derived cell populations, sites were further filtered using a rate of detection in at least 8 out of 9 samples per cell type. All remaining sites were annotated using ANNOVAR^51^ to gene symbols using RefGene and repeat regions using RepeatMasker v4.1.1^52^. Conservation metrics were gathered from the *phastConsElements30way* table of the UCSC Genome Browser, which consists of evolutionary conservation using *phastCons* and *phyloP* from the PHAST package^53^. Importantly, for sites uniquely detected in one cell type, we performed a third round of RNA editing quantification in the remaining two cell types to determine whether those sites in the neighboring cell types displayed high coverage and little-to-no editing (escaping the minimum coverage and edited read threshold) or simply little-to no read coverage (escaping the minimum coverage threshold). Resulting RNA editing data frames per cell type contained no more than ∼6% missing data. These values were imputed in a cell-specific manner using median imputation (i.e. taking the median editing rate across 8 donors per cell type). Resulting sites from these steps were subsequently referred to as high-confidence selective RNA editing sites and were used for downstream analysis.

To quantify RNA editing in snRNA-sesq and GTEx data, we called known sites from the REDIportal database in addition to any novel sites identified from FANS-derived cell populations. To this end, to quantify RNA editing sites and rates from a list of given sites, nucleotide coordinates for all such sites were used to extract reads from each sample using the samtools mpileup function, as previously described by us^17^. This approach quantifies the total number of edited reads and the total number of unedited reads that map to each RNA editing site detected. All analyses considered read strandedness when appropriate.

### Commonly used terms and definitions used in this study

*Cell type-specific RNA editing site*: Sites uniquely detected in one cell population and with zero coverage (or with insufficient detection rates) in the other two cellular populations.

*Cell type-enriched RNA editing site*: Sites detected in at least two cell population but with significantly higher RNA editing levels in one of the two cell types, determined by linear regression analysis.

### Computing the Alu Editing Index

The AEI method v1.0^54^ was leveraged to compute the Alu Editing Index for each sample using the STAR mapped bam files as input. The AEI is computed as the ratio of edited reads (A-to-G mismatches) over the total coverage of adenosines and is a robust measure that retains the full Alu editing signal, including editing events residing in low-coverage regions with a low false discovery rate.

### Quantification of RNA hyper-editing

RNA reads that undergo extensive hyper-editing of many neighboring adenosines within an extended region or cluster on the same transcript will not align to the reference genome due to the high degree of dissimilarity. Therefore, to identify hyper-edited reads in the current study, all unmapped reads from the original STAR alignment were converted to FASTQ and used as input for hyper-editing analysis. We adopted a well-established RNA hyper-editing pipeline^33^ with minor additional processing steps^15^, including: (1) extending cluster boundaries by the average distance between editing sites per cluster and subsequently merged clusters with overlapping coordinates (cluster length is a commonly product of read length); (2) all resulting hyper-editing sites were annotated using ANNOVAR (described above); (3) sites mapping to common genomic variation in dbSNP(v150) and those in gnomAD (maf>0.05) were discarded; (4) sites mapping to paired high-confidence private genomic calls were also discarded.

To minimize any batch effects, we computed a normalized hyper-editing signal per million mapped bases as previously described^33^. The normalized hyper-editing signal was computed by dividing the total number of resulting high-quality RNA hyper-editing sites over the total number of mapped bases from the STAR alignment. The number of total uniquely mapped bases for each sample were collected using Picard Tools v2.22.3 (http://broadinstitute.github.io/picard/) on each mapped bam file.

### Local motif enrichment analysis

EDLogo was used to quantify local sequence motifs^55^. We pulled sequences 4bp (±) the target adenosine to evaluate enrichment and depletion of specific nucleotides neighboring A-to-G RNA editing sites.

### Analysis of RNA editing by gene length

The total number of selective A-to-G RNA editing events were computed for each gene within each cell population. The total number of edits per gene were compared in a series of pairwise comparisons across each cell populations. To adjust for gene length, we residualized the number of A-to-G editing events per gene for each cell type by the log of gene length (geneEdits∼log(gene length)) using the resid function in the R package. Subsequent pairwise comparisons of the residualized number of RNA editing events per gene were performed (MGE-GABA vs. GLU, MGE-GABA vs OLIG, GLU vs. OLIG). Genes displaying an enrichment of residualized RNA editing sites were those dented as outlier genes beyond the 99% confidence intervals from the grand mean.

### RNA binding protein and eCLIP enrichment analysis

RNA binding proteins (RBPs) were first defined based on a consensus list of 837 high-confidence human RBPs from the hRBPome database^56^. A total of 470 RBPs were detected in FANS-derive cell populations and subjected to further analysis. Differential expression of these RBPs was computed using a linear model through the limma R package^57^ covarying for the possible influence of age and PMI. Donor as a repeated measure was controlled for using the duplicateCorrelation function in limma^57^. A matching approach was used to test for all genes genome-wide for subsequent analyses. To explore putative trans regulators of RNA editing, we further studied all RBPs that were significantly differentially expressed. RBPs were further subjected to correlation analysis with the AEI metric using a Pearson’s correlation coefficient.

Next, we collected transcriptome-wide binding patterns of FMRP (an RBP in the current analysis known to interact with *ADAR1* and *ADAR2*). Data from two eCLIP experiments and an input control experiment were obtained using postmortem frontal cortex from control subjects, as previously described^18^. The regioneR R package^58^ was used test overlaps of FMRP binding regions for each replicate separately with cell type-specific RNA editing sites based on permutation sampling. We repeatedly sampled random regions from the genome 1000 times, matching size and chromosomal distribution of the region set under study. By recomputing the overlap with FMRP binding sites in each permutation, statistical significance of the observed overlap was computed.

### Cell type-enrichment of RNA editing levels by differential editing analysis

To identify sites with differing levels of RNA editing between two given cell types, we implemented linear model though the *limma* R package^57^ covarying for the possible influence of age, PMI and donor as a repeated measure using the duplicateCorrelation function. Secondary models covaried for *ADAR1* and *ADAR2* expression (the enzymatically active editing enzymes) to explore potential ADAR-dependent editing activity. All significance values were adjusted for multiple testing using the Benjamini and Hochberg (BH) method to control the false discovery rate (FDR). Sites passing a multiple test corrected *P*-value < 0.05 were labeled significant.

### Gene ontology enrichment of differentially edited sites

Genes harboring differentially edited sites were functionally annotated using the ToppFunn module of ToppGene Suite software^59^. We set a genomic background defined as all genes harboring at least one editing site and tested for significance using a one-tailed hyper-geometric distribution with a Bonferroni correction. This is a proportion test that assumes a binomial distribution and independence for probability of any gene belonging to any set. We use a one-sided test because we are explicitly testing for over-representation of genes that harbor editing sites across hundreds of GO categories, without any *a priori* selection of candidate gene sets.

### Enrichment of cell-specific sites across neurodevelopment and neurological disorders

Both cell-specific and –enriched RNA editing sites were interrogated for over-representation of RNA editing sites previously found to be dysregulated across neurodevelopment and in neuropsychiatric disorders. Transcriptome-derived sets of RNA editing sites were curated based the following curation of RNA editing sites: two lists of sites found to change in editing rates across prenatal and postnatal cortical development^15, 16^; dysregulated RNA editing sites in schizophrenia^17^, Fragile X Syndrome and ASD^18^. To compute significance of all intersections, we used the GeneOverlap function in R which uses a Fisher’s exact test and an estimated odds-ratio for all pair-wise tests based on a background set of genes detected in the current study.

### Validation of recoding sites by targeted PCR amplification and high-throughput sequencing

We selected four cell type-specific recoding RNA editing events detected in our study to validate the results using an independent approach. We employed brain tissue samples from orbitofrontal cortex (OFC) in an independent cohort of 6 adult individuals without a neuropsychiatric diagnosis. Nuclei of MGE-GABA, GLU, and OLIG cells were isolated by FANS, followed by RNA extraction and library construction, as described above. PCR primers were designed to target four recoding RNA sites in: UNC80, GRIA3, GRIA4, and SON (**Figure S8, Supplemental Table 6**). The rhAmpSeq targeted amplicon sequencing kit (Integrated DNA Technologies) was used in two rounds of PCR amplification, to obtain the targeted PCR products (PCR1) and to introduce unique indexes for multiplex sequencing (PCR2), according rhAMPSeq protocol. Three µg of RNA-seq library was used in the first round of PCR for each subject and cell-type. The resulting rhAmpSeq libraries were cleaned using Agencourt AMPure XP beads, pooled, and sequenced on MiSeq obtaining at least 15,000 reads per sample (median read number per sample: 15,700 reads). The sequencing data (FASTQ files) were used to quantify the RNA editing read numbers and editing percentages at the four studied sites by counting reads which mapped identically to the +/-10 bases surrounded the edited site.

### snRNA-seq to identify and validate RNA editing sites

Mapped bam files were obtained following the alignment of short-reads to the reference genome (GRCh38/hg38). Reads were filtered for low quality, and counts were quantified (cell barcode counts and unique molecular identifier counts for each annotated gene) using CellRanger count, as previous described^46^. To optimize cell classification and reduce unwanted variance, we quality filtered, normalized, and scaled data according to Seurat’s guidelines, as previously described for this data set^46^. Like the original report, we identified 29 transcriptionally distinct cell clusters representing various populations of glutamatergic excitatory projection neurons, GABAergic interneurons, oligodendrocyte progenitor cells, oligodendrocytes, astrocytes, microglia, endothelial cells, and mural cells. Here, we further reduced these clusters into 24 cell clusters by collapsing five highly similar GLU cell subsets based on *NEUROD6* expression. Additionally, we further collapsed these cell clusters into six main cellular pools that constitute MGE-GABA, GLU and OLIG populations (based on the presence/absence of markers *RBFOX1*, *SOX6*, *SOX10*) and astrocyte (*GFAP*), microglia (*TNEN119*) and endothelial cell (*CLDN5*) populations.

To quantify RNA editing sites from snRNA-seq data, each mapped bam files was parsed into six unique bam files (per donor) reflective of the six main cellular pools (described above) using each cells unique molecular identifier (UMI). In this way, we pooled cells with matching cell type markers and gene expression patterns for subsequent RNA editing analysis. Next, we quantified RNA editing levels for all high-confidence sites identified in the FANS-derived cell populations using the samtools mpileup function (described above). Given the challenges related to the limited number of cells per pool and low sequencing depth, we explored validation rates under varying read coverage thresholds (minimum of 3-20 reads), but ultimately required a minimum coverage of 10 reads per site, as applied for all other sites in the current study.

### GTEx data pre-processing and cell type deconvolution

All GTEx FASTQ files were mapped to human reference genome (GRCh38) using STAR v2.7.3^47^ and counted using featureCounts^60^. Count matrices were assembled for each brain region, filtered to retain genes with at least 1 count per million in at least half of samples per region, and VOOM normalized using limma^57^. Each resulting normalized data frame was subjected to principal component analysis to identify and remove any outlier samples that lay beyond two standard deviations from the grand mean. Following outlier removed, a total of 1,129 samples were retained for all subsequent analyses.

To compute cellular composition, we applied non-negative least squares (NNLS) from the MIND R package^61^ and utilized the Darmanis et al., signature matrix^62^ which contained a mixture of six major cell types: astrocytes, oligodendrocytes, microglia, endothelial cells, excitatory and inhibitory neurons. NNLS, executed through the est_frac function, was applied to log_2_ count per-million (CPM) transformed data using the *limma* package in R. For each sample, both excitatory and inhibitory neuronal predictions were summed into one ‘neuronal’ cell population. We focus our predictions on these major cell types in an effort to reduce noise and to evaluate a distribution of cell types that reflect an approximate expected distribution in the human brain based on prior work.

### GTEx bulk brain cis-edQTL analysis

We conducted *cis*-eQTL mapping within the 13 brain regions. We leveraged existing VCFs for high quality imputed genotype array data from dbGAP (phg000520.v2.GTEx.MidPoint.Imputation.genotype-calls-vcf.c1.GRU.tar), using SNPs with an and estimated minor allele frequency ≥ 0.05. Only RNA editing sites detected in at least 50% of samples per region were considered for *cis*-edQTL mapping. Each data matrix for each region exhibited ∼17% missing values, which were imputed using the well-validated predictive mean matching method in the *mice* R package using 5 multiple imputations and 30 iterations^63^. The genomic coordinates for all RNA editing sites were lifted over to GRCh37. Subsequently, to map genome-wide edQTLs, a linear model was used on the imputed genotype dosages and RNA editing levels using MatrixEQTL^64^. RNA editing levels were covaried for sex, age, RIN and type of death. To control for multiple tests, the FDR was estimated for all *cis*-edQTLs (defined as 1Mb between SNP marker and editing position), controlling for FDR across all chromosomes. Significant *cis*-edQTLs were identified using a genome-wide significance threshold (FDR < 0.05). To assess whether *cis*-edQTLs relate to brain promoter and enhancer regions, overlap between max cis-edQTLs and promoter and enhancer regions was tested for from the FANTOM project collected from the SlideBase database^65^. A permutation-based approach with 1,000 random permutations was used to determine statistical significance of the overlap between edSNP coordinates and enhancer regions using the R package regioneR^58^.

### Data and code availability

To promote the exchange of this information, we developed an interactive Rshiny app with an easily searchable interface to act as a companion site for this paper: https://breenms.shinyapps.io/CNS_RNA_Editing/. This application allows users to download sites along a specific gene of interest. Additionally, all code, major data tables and matrices as well as summary statistics are provided in GitHub: https://github.com/BreenMS/RNA-editing-in-CNS-cell-types and https://github.com/ryncuddleston/RNA-hyper-editing.

## Author contributions

Conceptualization: MSB, SD, EAM

Data analysis and interpretation: MSB, RC, JL, SK, XF, ML, EAM

Provided revisions to scientific content: ML, SK, AK

Visualization: MSB, RC, XF

Funding acquisition: MSB, SD

Supervision: MSB, SD

Writing –original draft: MSB, RC, JL, XF, AK, ML, SD, EAM

## SUPPLEMENTAL FIGURES

**Figure S1.**
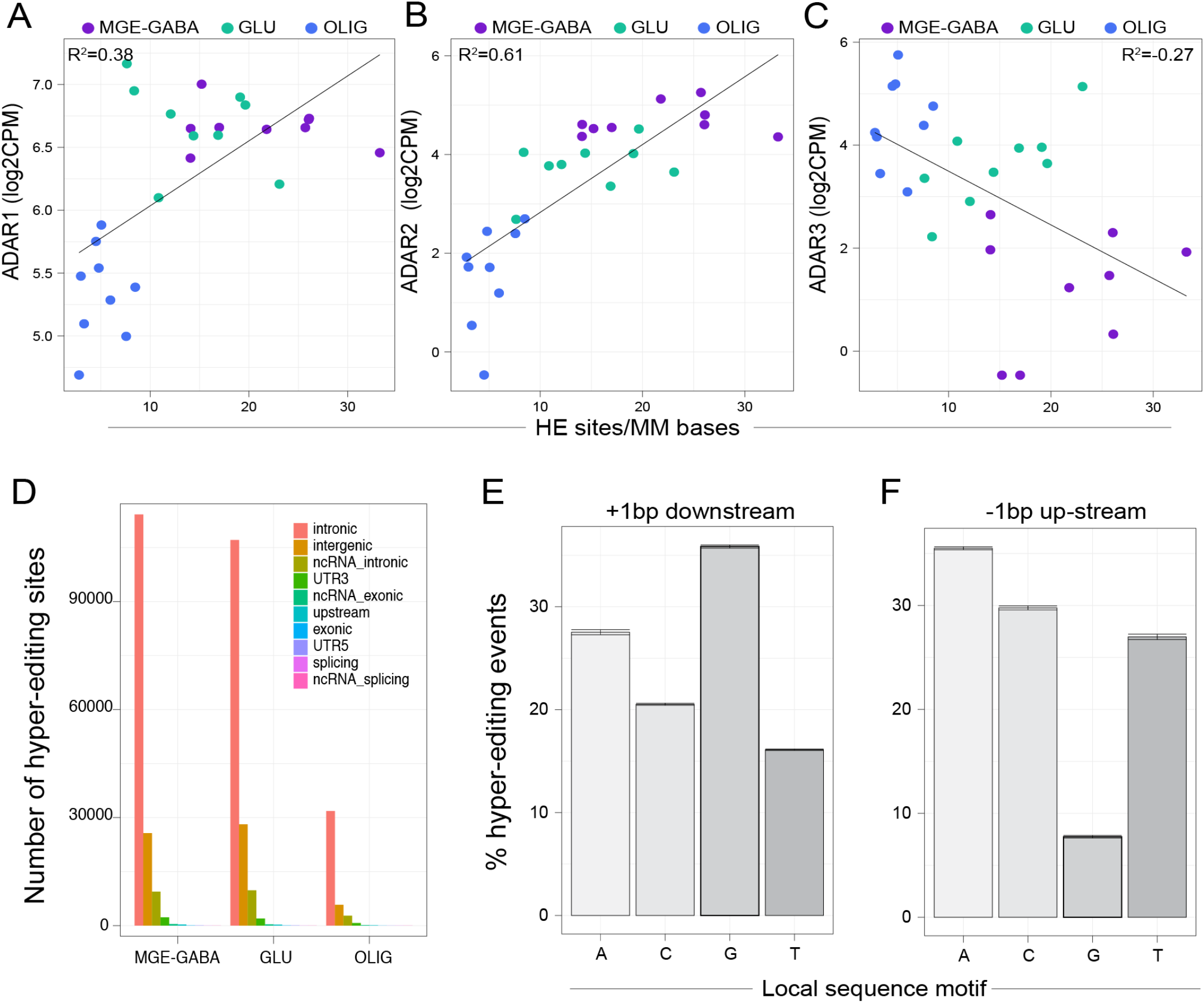
RNA hyper-editing across cell types. The amount of normalized hyper-editing per million mapped bases (HE/MM, x-axis) variance explained by expression of (**A**) *ADAR1*, (**B**) *ADAR2*, and (**C**) *ADAR3* was evaluated by linear regression. (**D**) The total number of hyper-editing sites (y-axis) by genic region per cell type (x-axis). Hyper-editing sites were scrutinized for conserved local sequence motifs, that is (**E**) enrichment of guanosine +1 bp downstream (**F)** and depletion of guanosine −1 bp upstream of the target adenosine. Values are aggregated across all three cell types.

**Figure S2.**
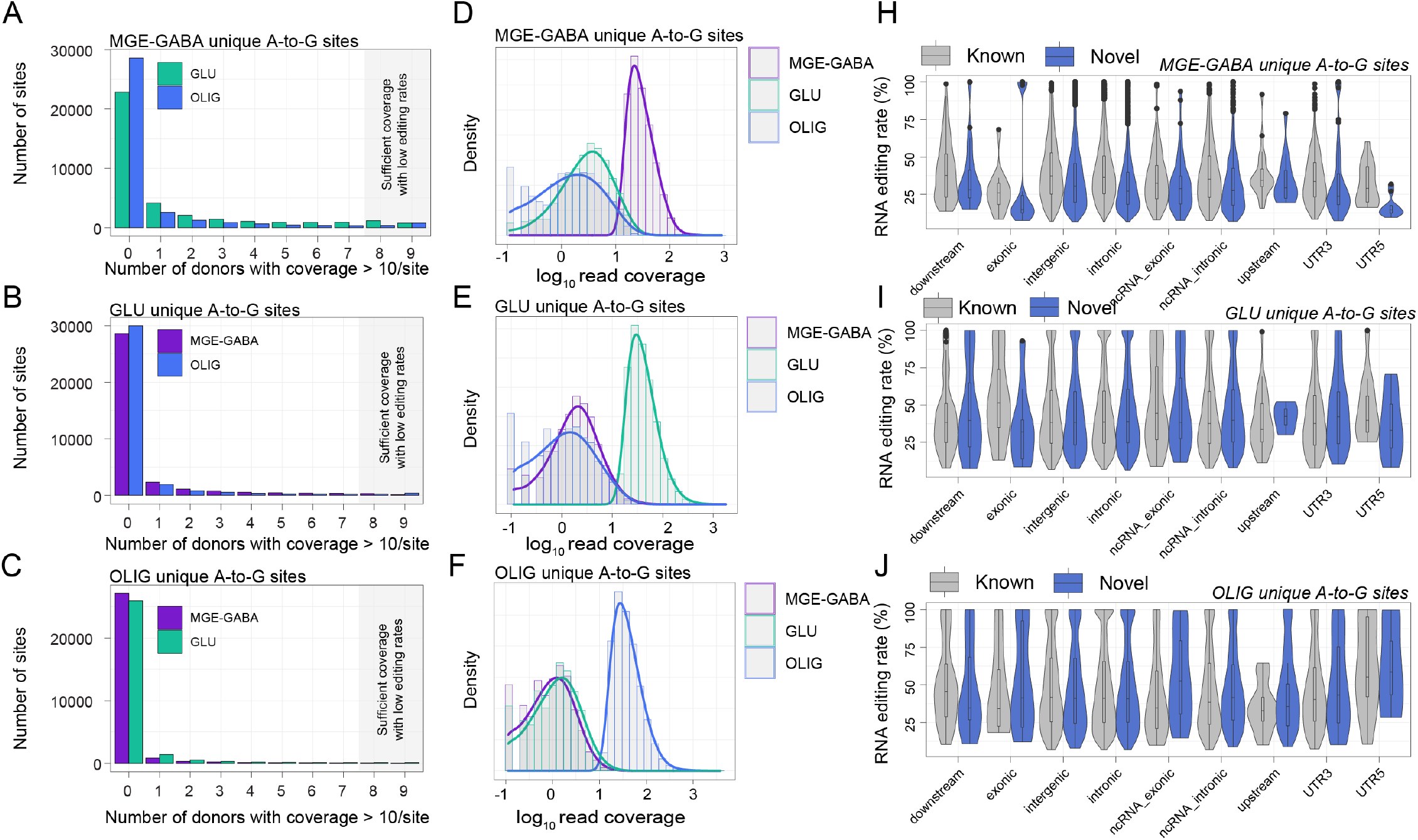
Annotation of A-to-G sites detected in only one cell type. The number of donors (out of 9 per cell type) with a coverage of at least 10 reads for sites uniquely classified as cell type-specific in **(A)** MGE-GABA, **(B)** GLU, and **(C)** OLIG cells. Read coverage plots for sites uniquely classified as cell type-specific in **(D)** MGE-GABA, **(E)** GLU, and **(F)** OLIG cells. Plots demonstrate that sites uniquely detected per each cell type are largely associated with differences in cell type-associated read coverage differences. RNA editing levels for both known and novel A-to-G cell-specific sites parsed by genic region for **(H)** MGE-GABA, **(I)** GLU, and **(J)** OLIG cells.

**Figure S3.**
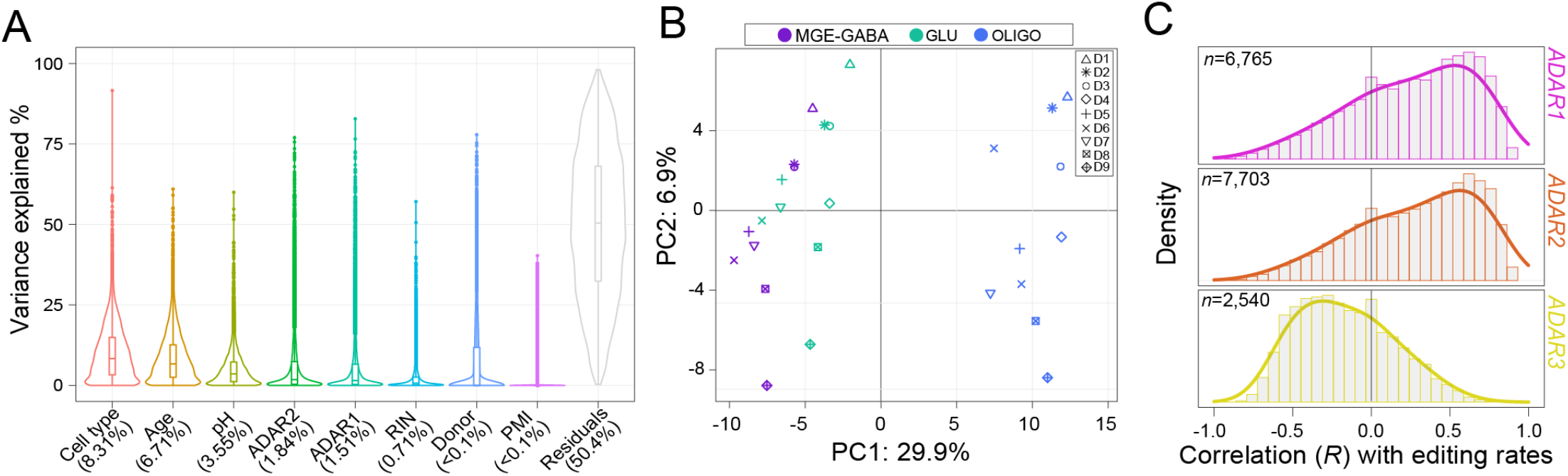
Quantifying variance in RNA editing rates by known factors. Sites commonly detected across all cell types were leveraged for the analyses presented in all current panels. (*n*=15,221 sites). **(A)** Mixed linear models were used to compute variance explained (%, y-axis) by several known factors and FANS-derived cell population (cell type) explained the largest source of variance (8.31%). **(B)** Principal component analysis of all donors (*n* = 9) stratifies neuronal (MGE-GABA, GLU) from non-neuronal (OLIG) cells. Samples are shaped differently based on donor (D) as a repeated measure. **(C)** Density plot distributions illustrate the number of A-to-G editing sites which correlate with ADAR expression across all cell-types. The total number of sites that are significantly associated (FDR P-value < 0.05) with ADAR expression are displayed in the top left corner for each enzyme.

**Figure S4.**
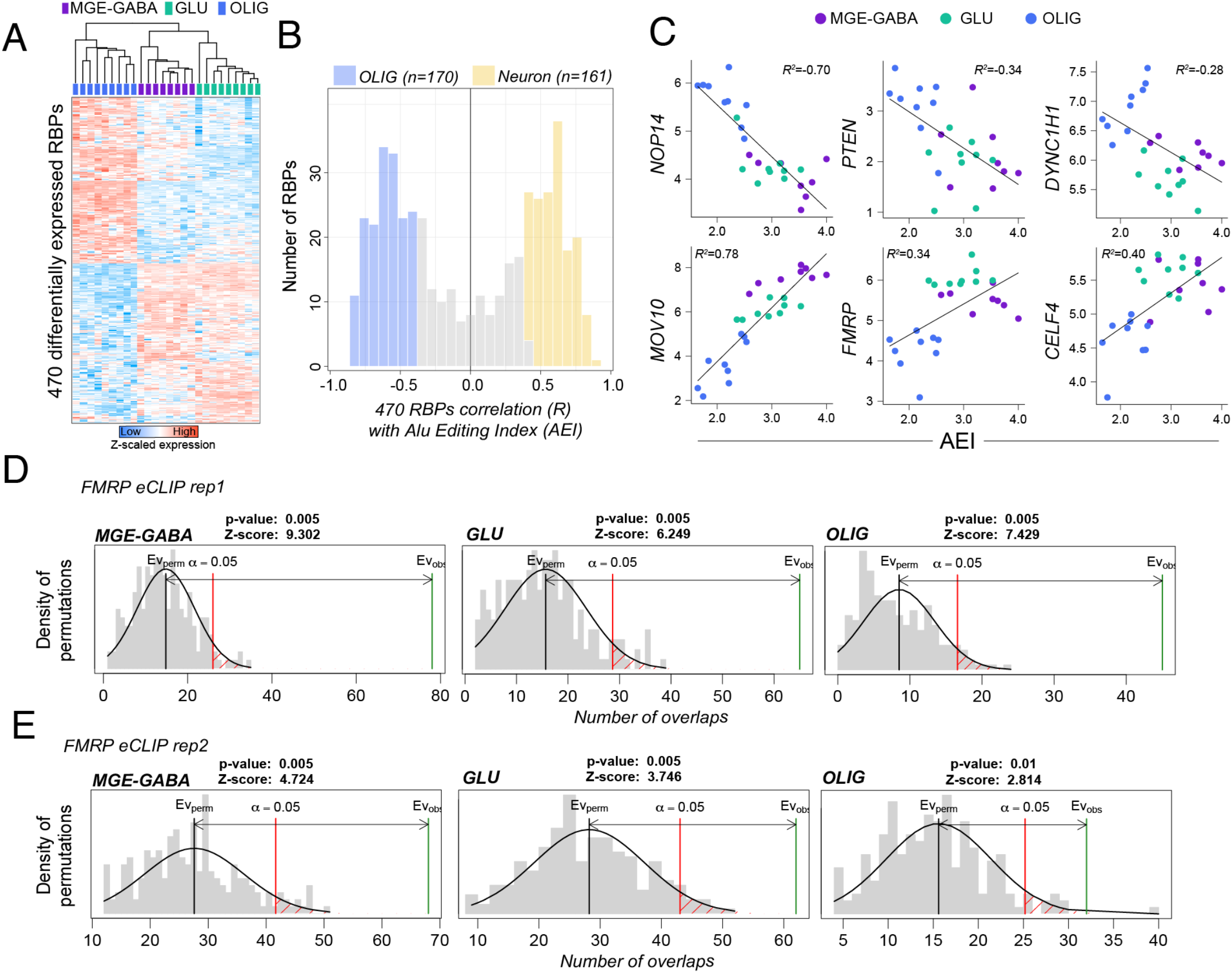
RNA editing variation and RNA binding protein analysis. **(A)** Heatmap displaying the relative expression of 470 differentially expressed RNA binding proteins (RBPs) between MGE-GABA (purple) and GLU (green) neurons relative to OLIG (blue) cells. **(B)** Correlation of 470 RBPs with the AEI measurement across all cell types depicting a 161 RBPs strongly associated with the AEI in neuronal cells (MGE-GABA and GLU) and 170 RBPs strongly associated with the AEI in OLIG cells. Analyses were adjusted for repeated measures using Benjamini-Hochberg method. (**C**) Example of three RBPs strongly associated with the AEI in OLIG cells (top) and three RBPs strongly associated with the AEI in neuronal cells (bottom). Genomic coordinates for all cell type-associated RNA editing sites were assessed for enrichment for FMRP eCLIP binding sites across two technical replicates (**D-E**). The regioneR R package was used test overlaps of genomic regions based on permutation sampling. We repeatedly sampled random regions from the genome1000 times, matching size and chromosomal distribution of the region set under study. By recomputing the overlap with FMRP binding sites in each permutation, statistical significance of the observed overlap was computed.

**Figure S5.**
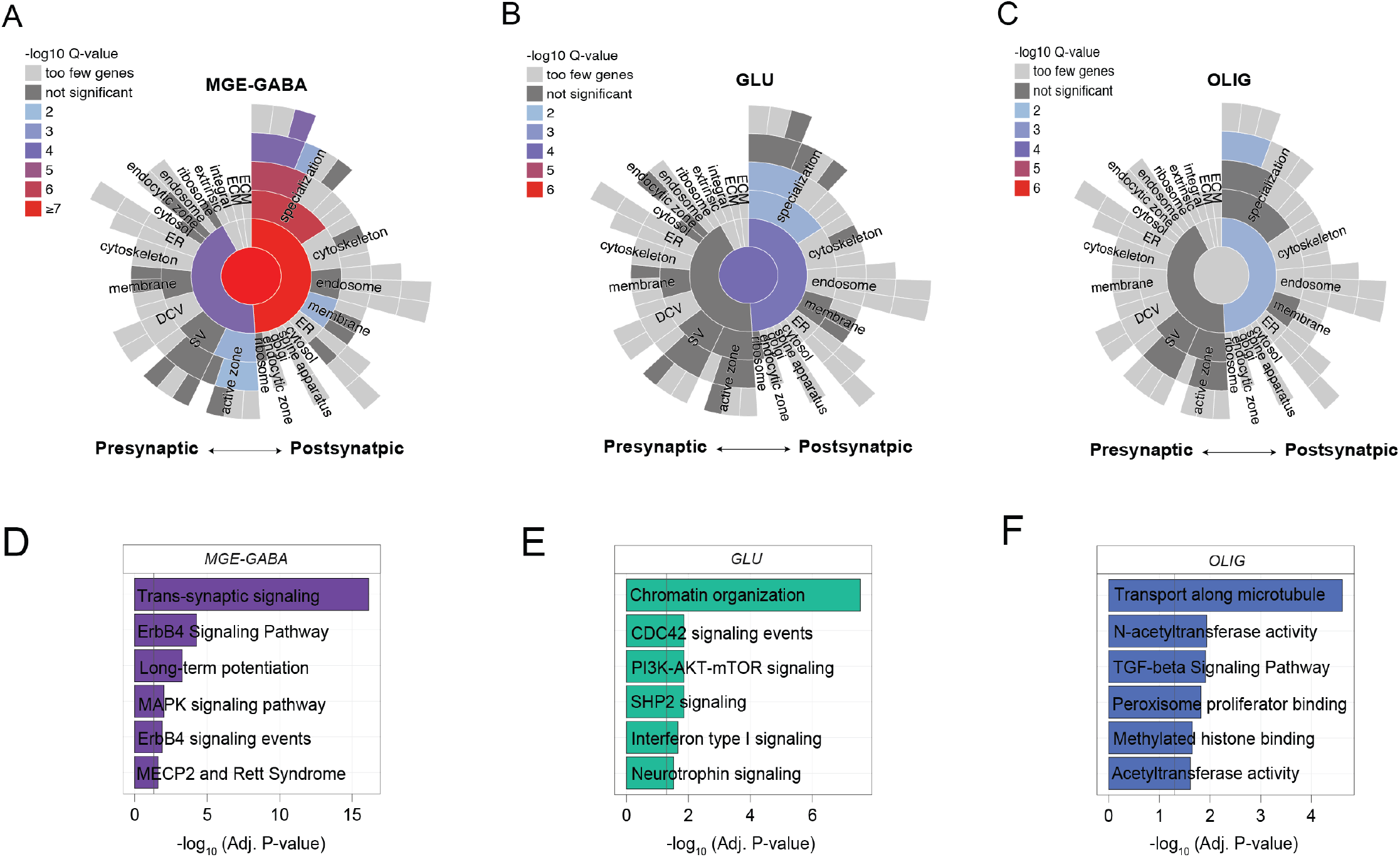
Synaptic gene-set enrichment analysis. SynGO sunburst plots portray synaptic gene ontology enrichment associated with cell-type enriched RNA editing sites in **(A)** MGE-GABA, **(B)** GLU, and **(C)** OLIG cells. The top six cell-specific pathways (i.e. uniquely enriched for one cell type only) for the genes encompassing identified editing sites for in **(D)** MGE-GABA, **(E)** GLU, and **(F)** OLIG cells.

**Figure S6.**
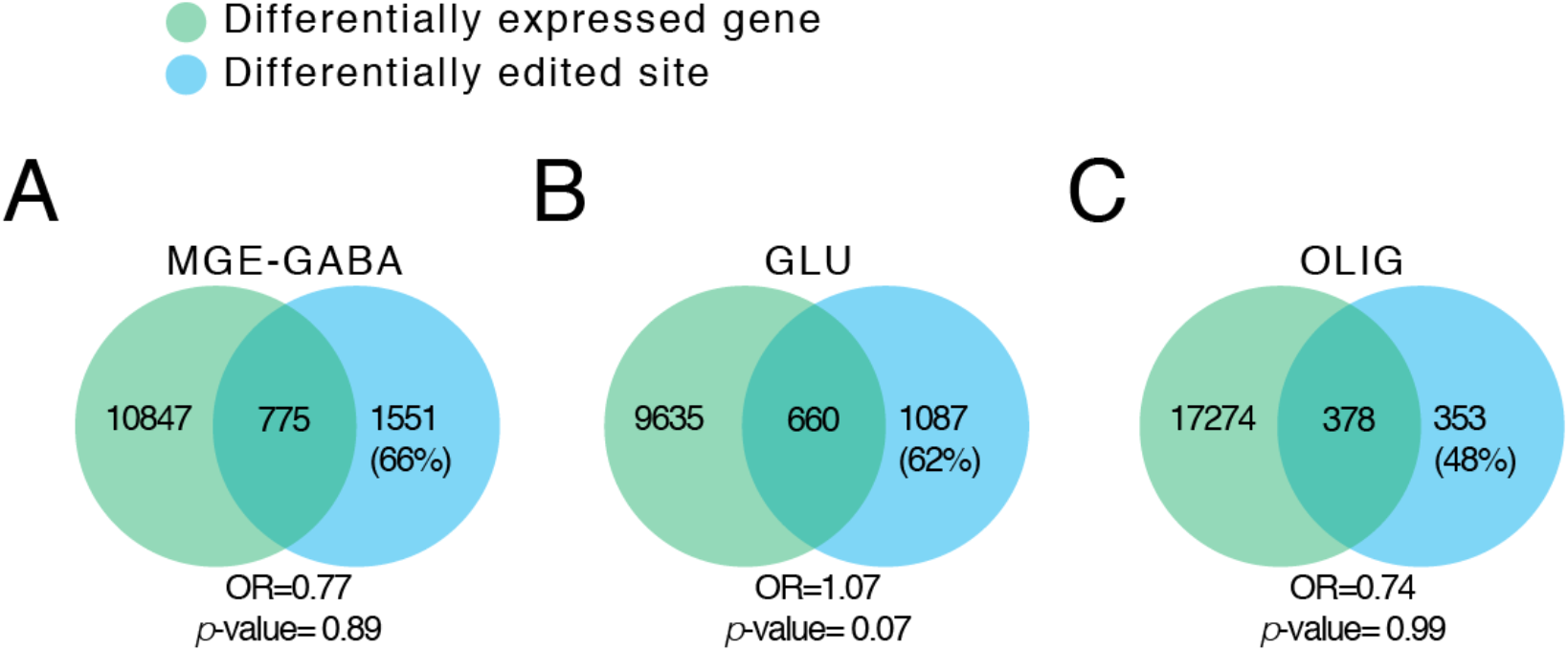
Overlap of differentially edited sites and differentially expressed genes. Cell-specific gene expression was defined as genes with significantly higher expression (FDR P-value < 0.05) in one cell type relative to other cell types. A Fisher’s Exact Test and an estimated odds-ratio (OR) was used to compute the significance of the overlap between cell-specific gene expression and genes harboring editing sites with cell-specific editing rates. We observed (**A**) ∼66% of differentially edited sites in MGE-GABA nuclei were not explained by differential gene expression, (**B**) ∼62% of differentially edited sites in GLU nuclei were not explained by differential gene expression, and (**C**) ∼48% of differentially edited sites in OLIG nuclei were not explained by differential gene expression.

**Figure S7.**
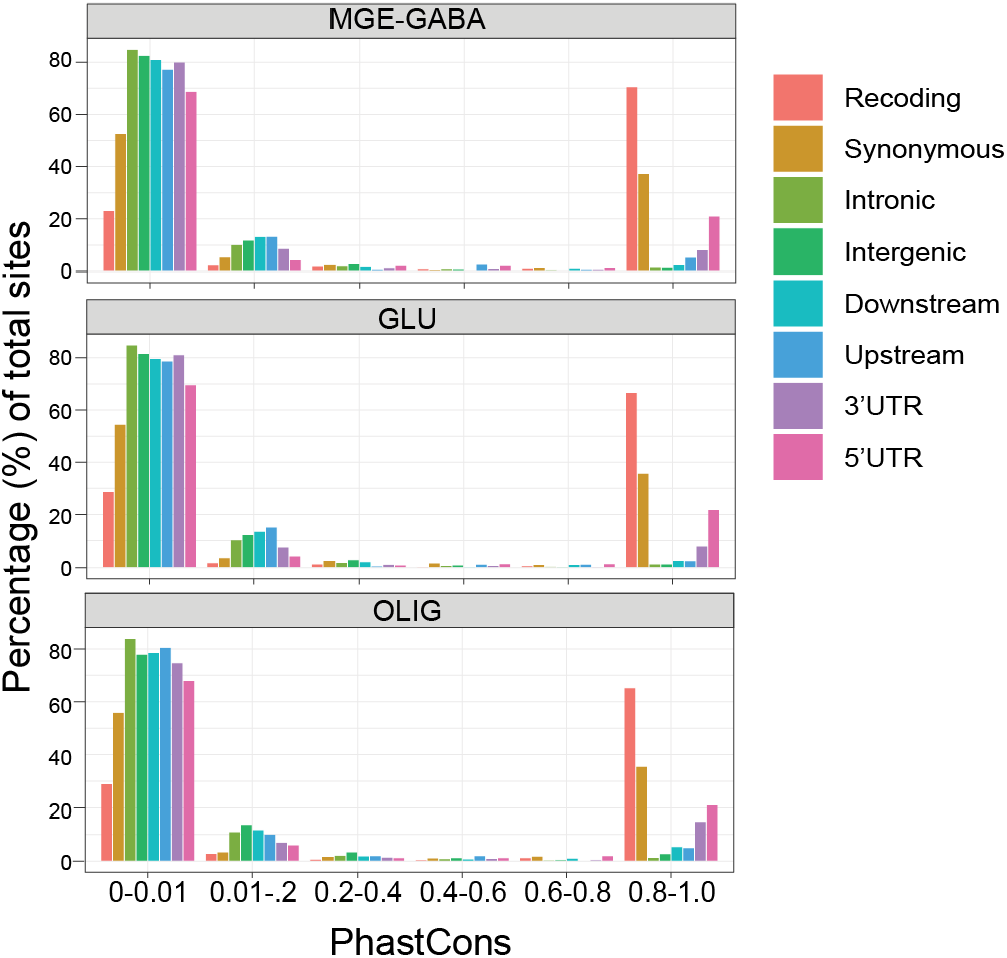
PhastCons scores by genic regions. Editing sites reported for all three cell types binned by genic region and PhastCons, highlighting an increased proportion of highly conserved recoding sites across all three cell types, with less conservation in editing sites throughout other genic regions.

**Figure S8.**
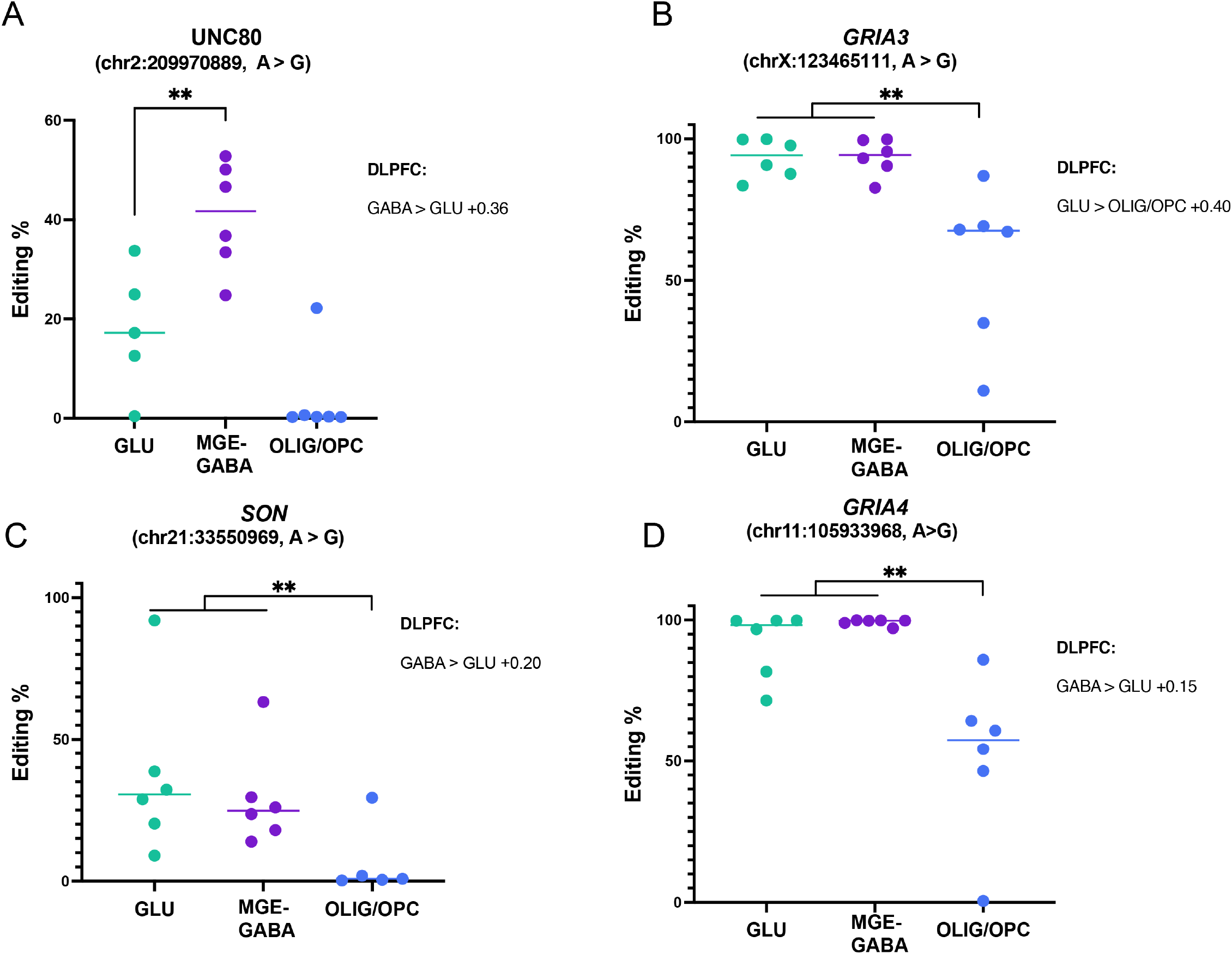
Validation of cell-specific recoding events in independent samples. Four recoding sites were validated in six independent OFC samples from healthy age matched donors.

**Figure S9.**
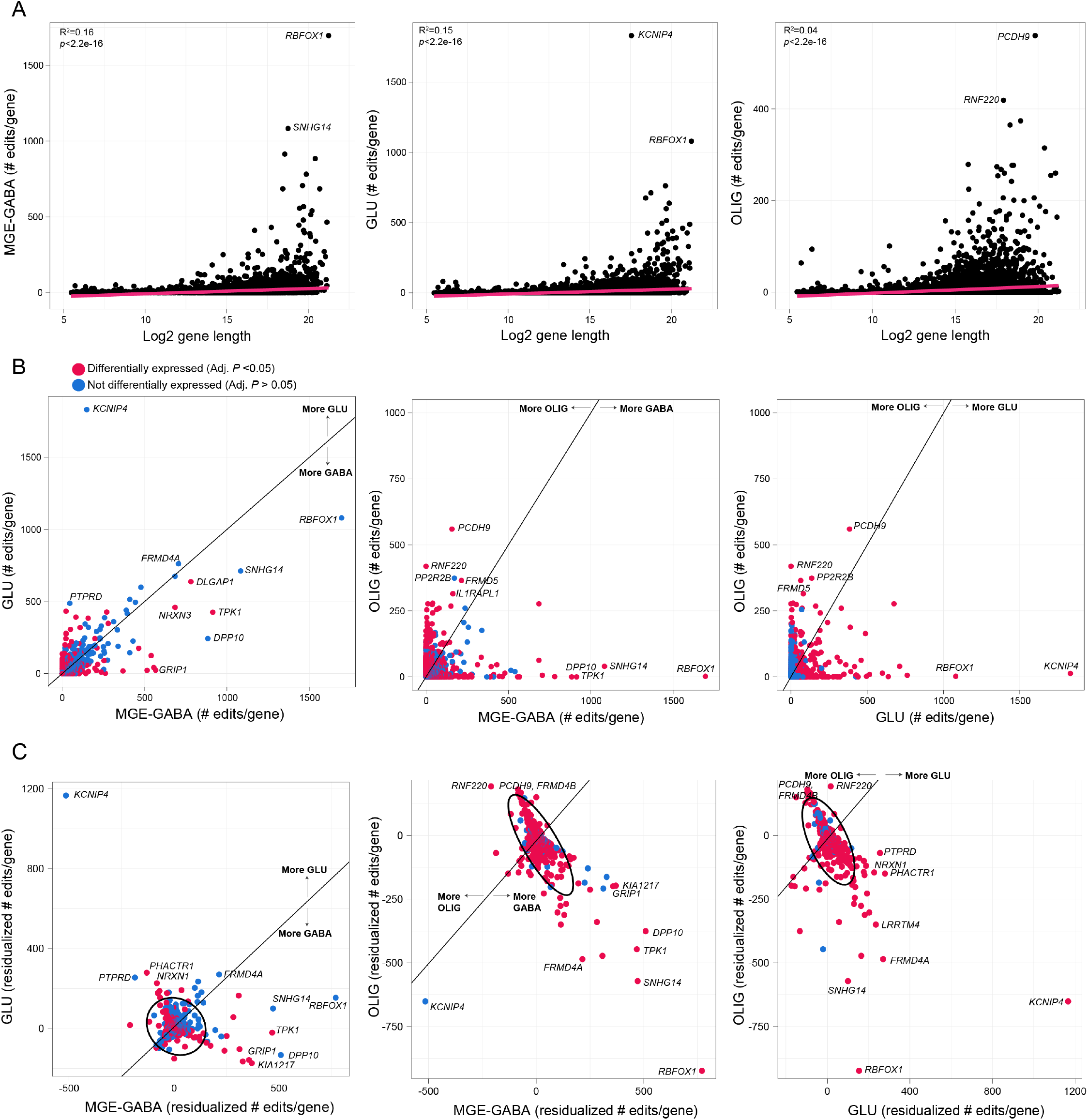
RNA editing sites as a function of gene length. **(A)** Gene length as a function of the total number of selective RNA editing sites across all MGE-GABA (left), GLU (center) and OLIG cells (right). **(B)** Pairwise comparisons of concordance for the number of RNA editing sites per gene between two cell types – MGE-GABA vs. GLU (left), MGE-GABA vs OLIG (center), GLU vs. OLIG (right). **(C)** Pairwise comparisons of concordance for the number of residualized RNA editing sites by gene length per gene between two cell types – MGE-GABA vs. GLU (left), MGE-GABA vs OLIG (center), GLU vs. OLIG (right). Outlier genes were classified as genes enriched with RNA editing sites and are denoted as those outside the 99% confidence intervals from the grand mean.

**Figure S10.**
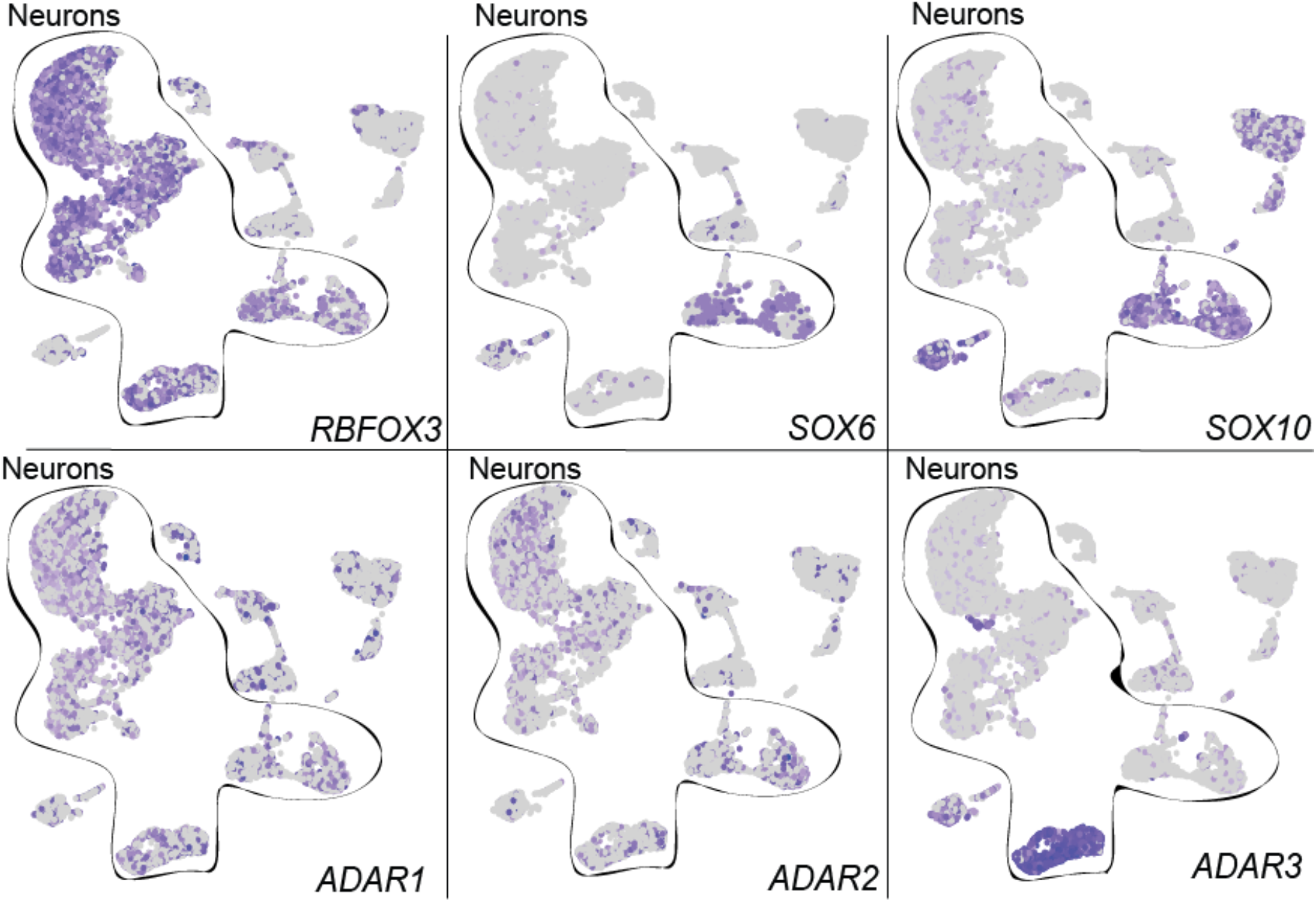
tSNE of snRNA-seq. Cells were examined for expression markers: GLU (RBFOX3+/SOX6-), MGE-GABA (RBFOX3+/SOX6+), and OLIG (RBFOX3-/SOX10+) (top row) and *ADAR1*, *ADAR2* and *ADAR3* expression (bottom row).

**Figure S11.**
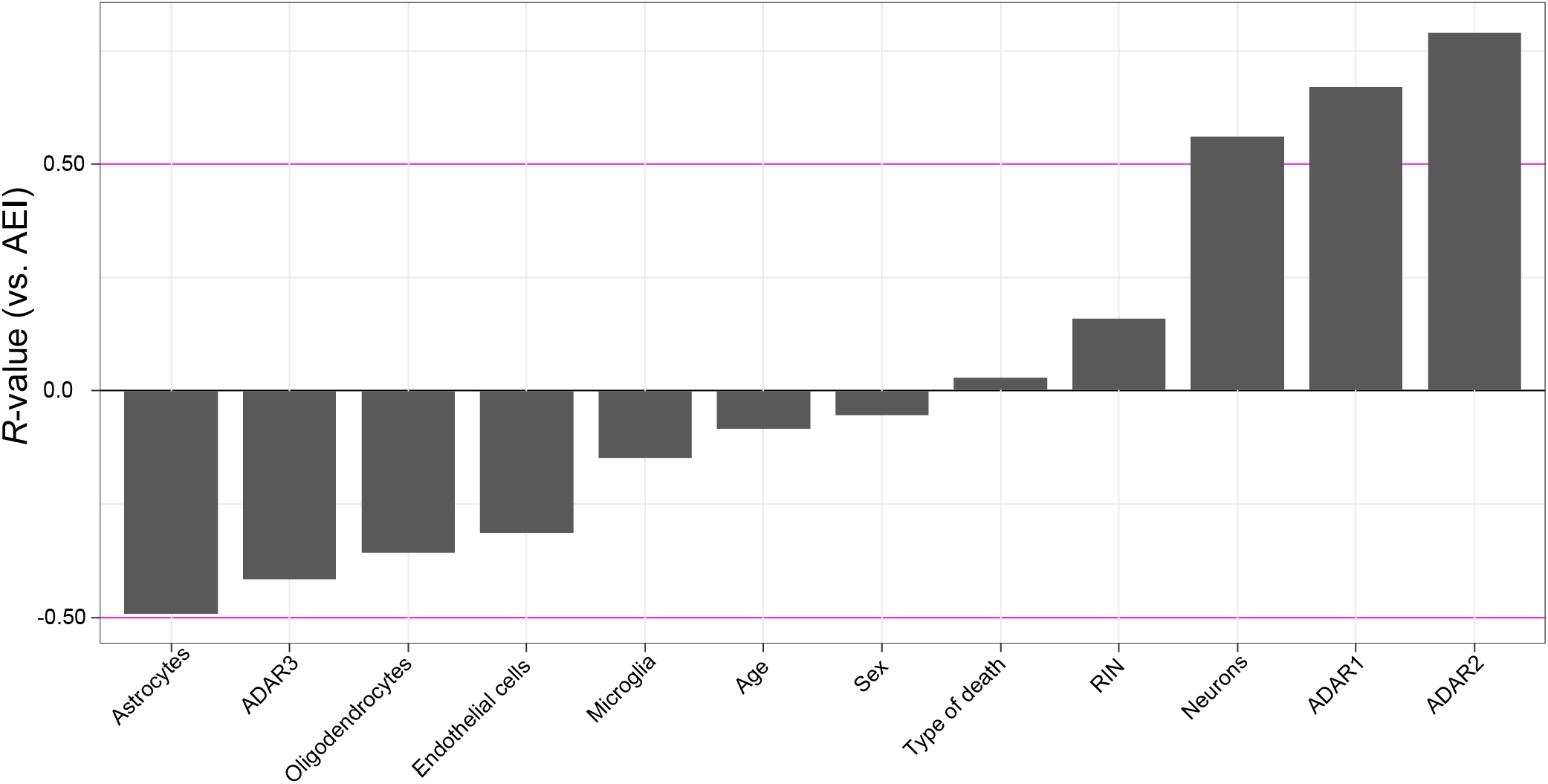
Correlation of AEI with known factors in GTEx data. The AEI was correlated with 12 biological factors, including cell type proportions, using a Pearson’s correlation. Correlations were performed across all regions. *ADAR2*, *ADAR1* and neuronal proportions display the strongest positive associations with AEI where as non-neuronal cell types and *ADAR3* display the strongest negative associations with the AEI.

**Figure S12.**
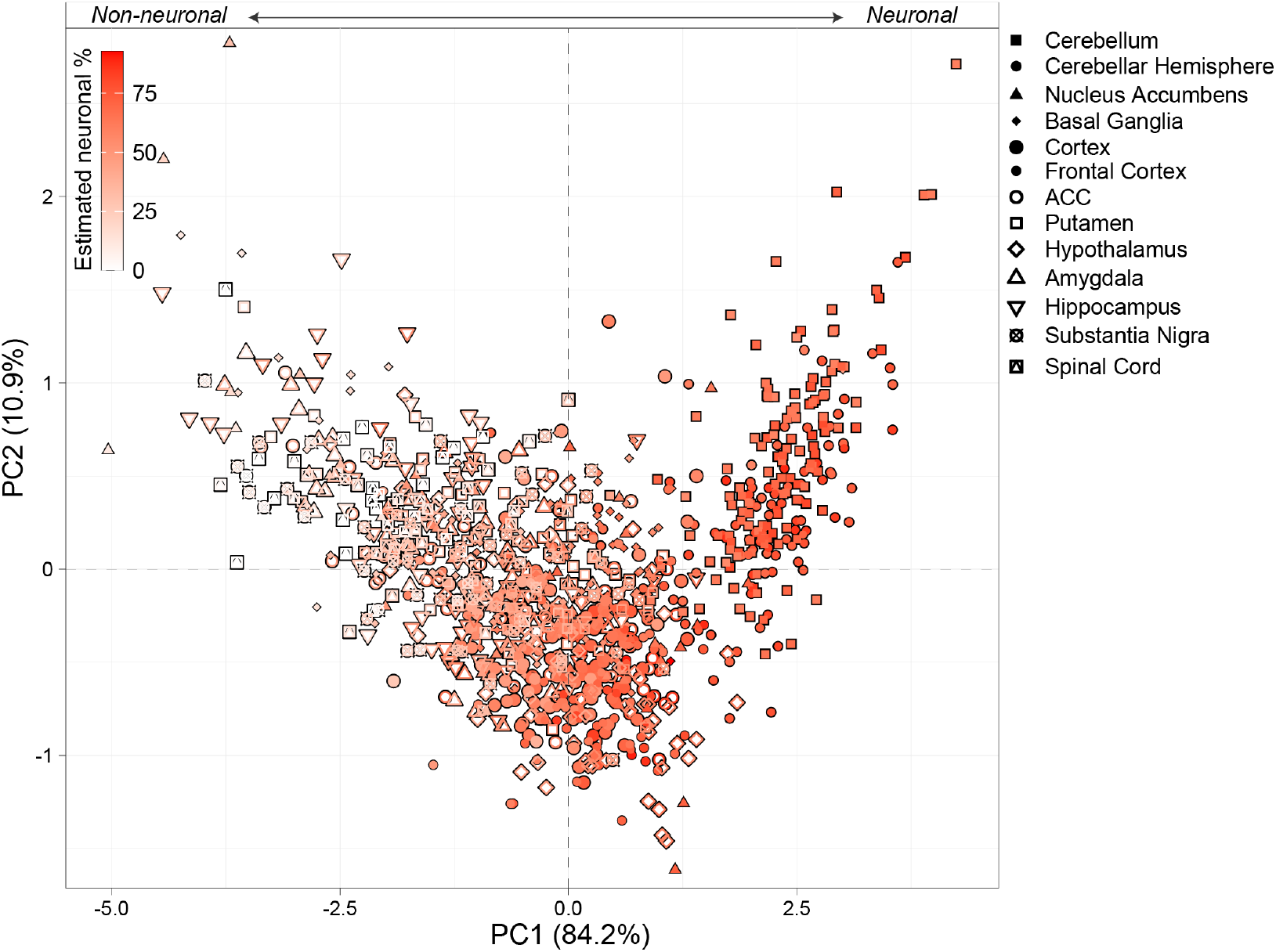
Principal component analysis on features of RNA editing in the brain. We performed PCA on ADAR1, ADAR2 and the AEI for each sample across all brain regions. We then color coded each sample by the proportions of neurons per donor, from red-(high, more neuronal) to-white (low, less neuronal). This result indicates that PC1 (based on features of RNA editing in bulk brain tissues) is largely associated with differences in neuronal proportions.

**Figure S13.**
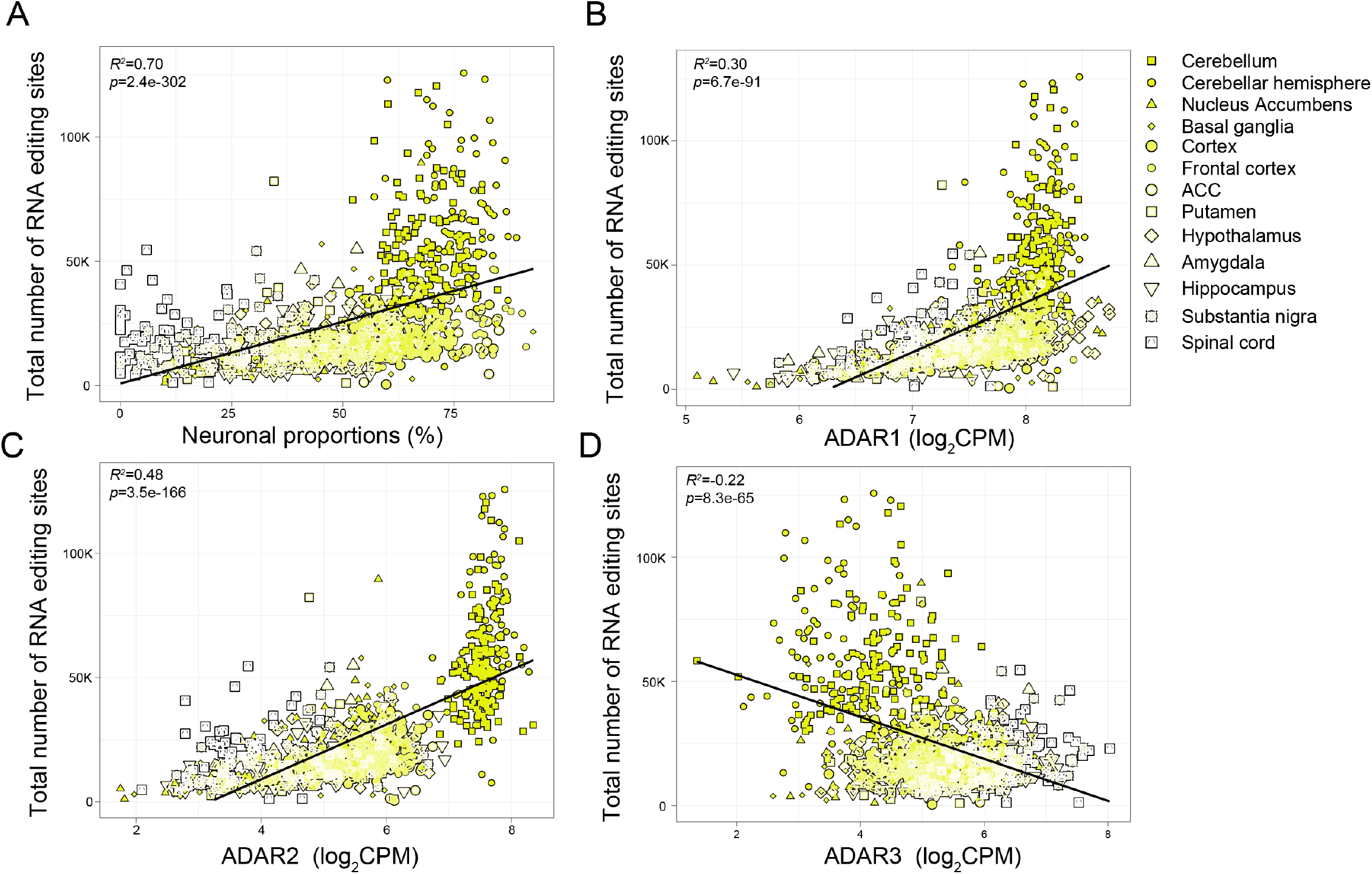
Variance in RNA editing site detection per donor in bulk GTEx tissues. The amount of variance explained in the total number of RNA editing sites detected (y-axis) across all brain regions (denoted by point shape) according to differences in (**A**) neuronal proportions, (**B**) *ADAR1*, (**C**), *ADAR2*, (**D**) and *ADAR3* expression. Linear regression was used to test the strength of significance for each association.

**Figure S14.**
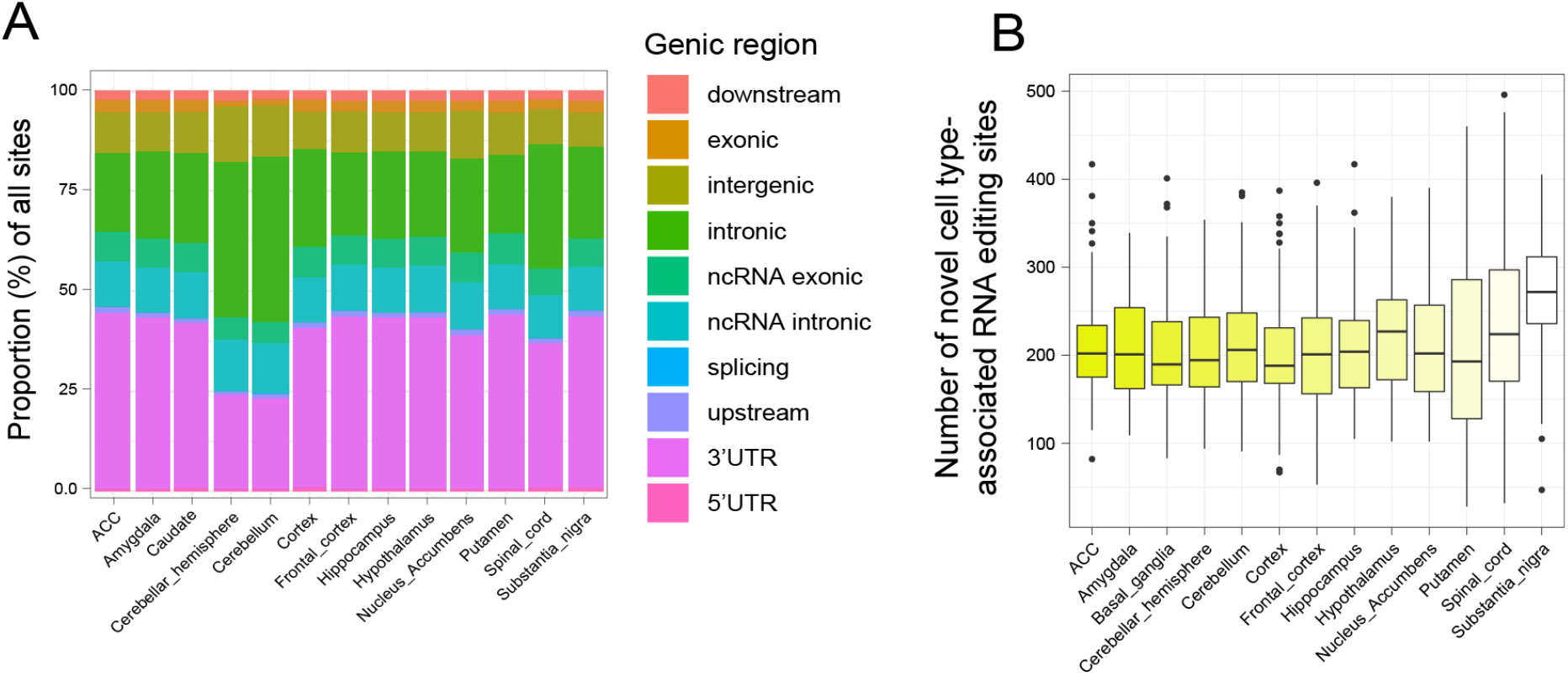
Annotation of sites detected in bulk GTEx brain tissue. (**A**) The fraction of RNA editing sites per genic region according to brain regional differences. (**B**) The number of novel cell type-associated RNA editing sites detected in bulk brain RNA-sequencing data across all regions (∼213 novel sites per region were detected).

**Figure S15.**
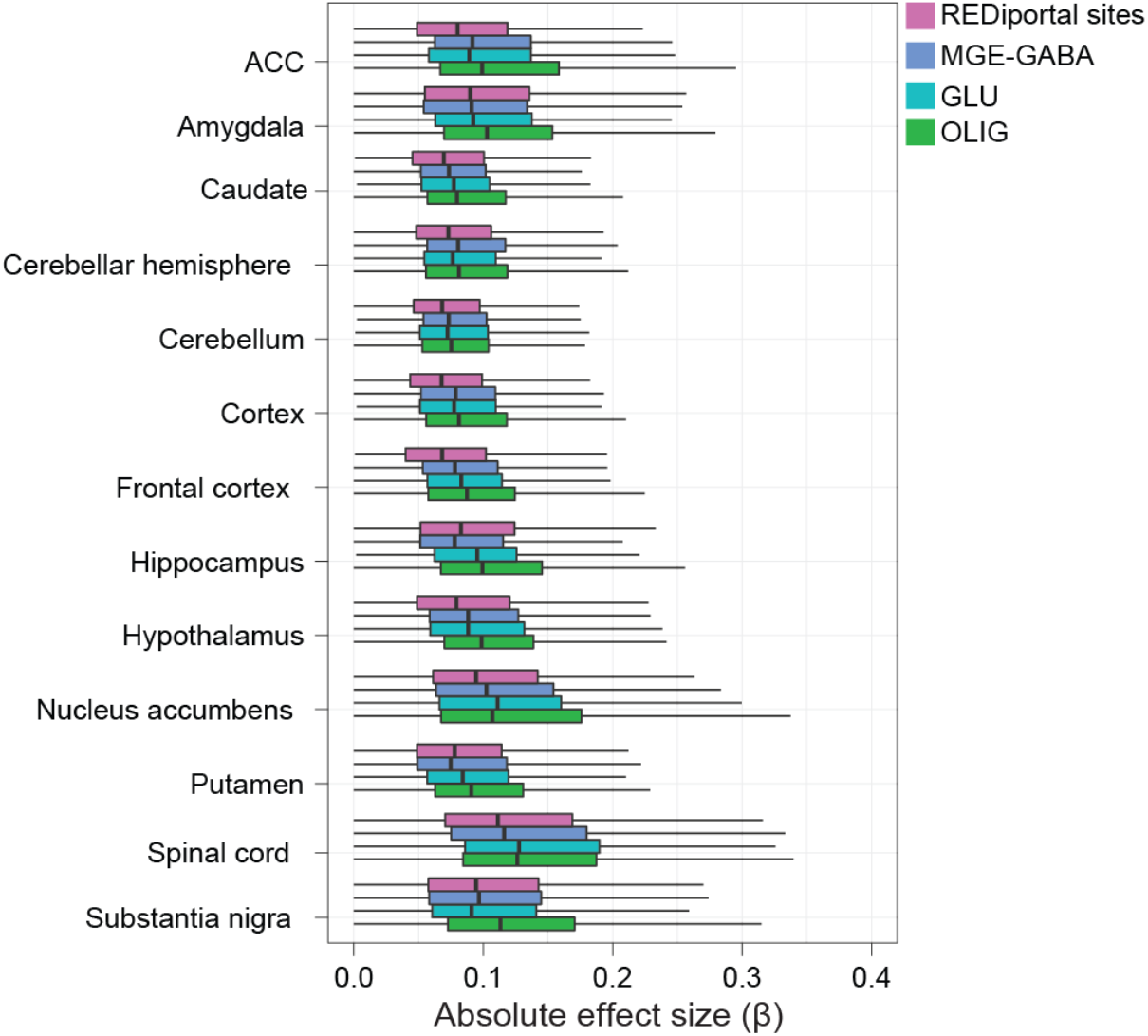
Absolute effect sizes of edQTLs across GTEx brain regions. Independent boxplots for each brain region express that the absolute effect sizes (β) are consistently higher for edQTLs related to cell-associated editing sites and lower for all other edQTLs for editing sites without cellular annotation.

**Figure S16.**
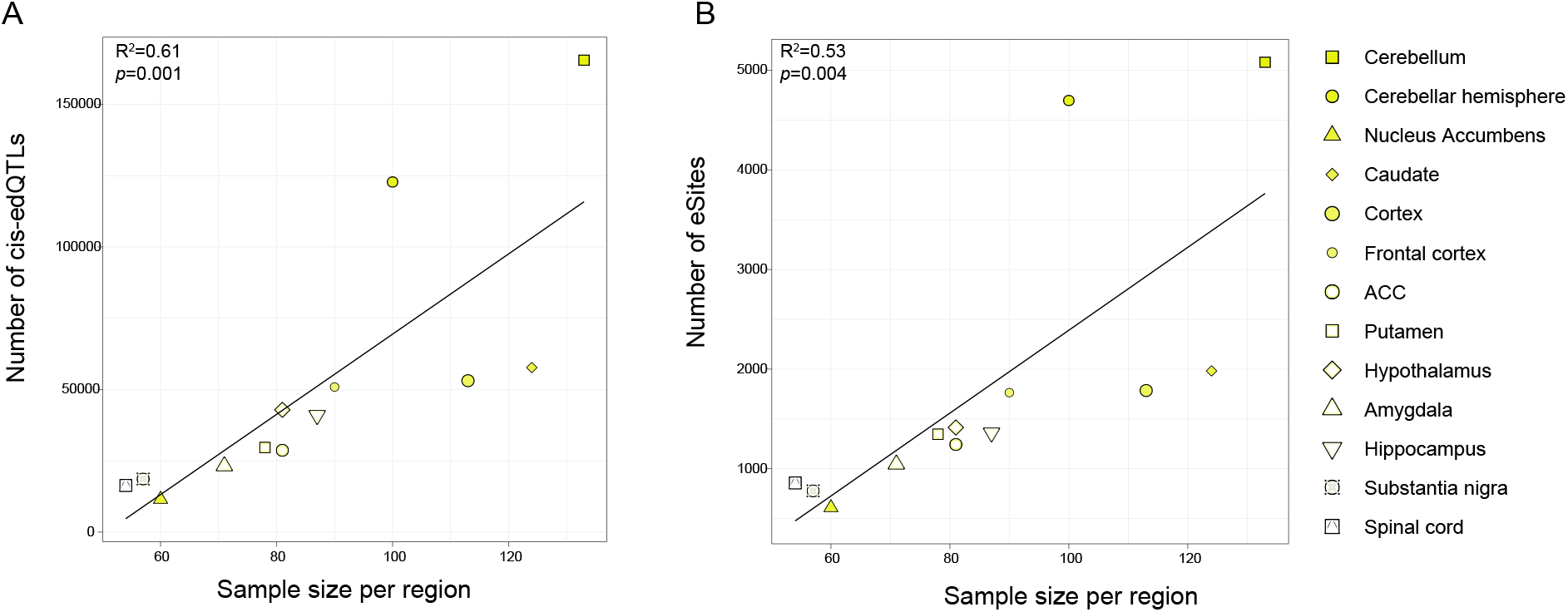
edQTL discovery as a function of sample size. **(A)** cis-edQTL **(B)** and eSite discovery as a function of sample size per brain region.

## Supplemental Tables

**Table S1.** Alu editing index and hyper-editing across purified cortical cell types.

**Table S2.** Annotation of cell-specific RNA editing sites.

**Table S3.** RNA editing sites as a function of gene length.

**Table S4.** RNA editing sites associated with ADAR and RBP expression.

**Table S5.** Differential RNA editing across cell types.

**Table S6.** Sample metadata and primer information for rhAmpSeq validation of recoding RNA-editing sites.

**Table S7.** RNA editing recoding events.

**Table S8.** snRNA-seq cellular pools.

**Table S9.** Features of global editing across bulk GTEx brain regions.

**Table S10.** Enrichment of cell-specific sites across GTEx brain regions.

**Table S11.** Max-edQTLs across bulk GTEx brain regions.

